# Genome-resolved metagenomics reveals a phylogenetically cohesive *Acetilactobacillus*-like species complex dominating stingless bee pot honey

**DOI:** 10.1101/2025.06.25.661387

**Authors:** Aurora Xolalpa-Aroche, Haydeé Contreras-Peruyero, Enrique J. Delgado-Suárez, David I. Hernández-Mena, Wilson I. Moguel-Chin, J. Fausto Rivero-Cruz, Rodrigo A. Velarde, Elizabeth Ortiz-Vázquez, Blanca E. Rivero-Cruz, Jose Abel Lovaco Flores, Lorena Rodríguez Orduña, Cuauhtémoc Licona Cassani, Francisco Barona-Gómez, Nelly Sélem-Mojica

## Abstract

Pot honey, the honey produced by stingless bees, is valued for its antimicrobial capacity, which may be influenced by its microbial content. While Lactobacillaceae species are commonly associated with honeybees and honey microbiomes, most studies have focused on *Apis mellifera*, leaving pot honey microbial diversity largely unexplored. We present the first pot honey shotgun metagenomic analysis from bee species *Melipona beecheii* and *Scaptotrigona mexicana*. We reconstructed 24 metagenome-assembled genomes (MAGs), 15 of which lacked close matches to any described species, showing *≤*81% Average Nucleotide Identity (ANI) to available reference genomes. Phylogenetic analyses resolved these MAGs into four well-defined clades (intraclade ANI *>* 99%, interclade ANI *≤* 81%), consistent with four novel species within the family Lactobacillaceae. GTDB-Tk classification placed MAG clades 1 and 2 closest to *Nicoliella*, and clades 3 and 4 closest to *Acetilactobacillus*. We validated the presence of these lineages in honey by sequencing three isolates that clustered within MAG clade 2. Aminoacid similarity (AAI/cAAI) indicates the presence of two genus-level lineages: one occupying a transitional genomic space near *Nicoliella*, and a second representing an undescribed genus. The genomic similarity of our MAGs and isolates to those from pot honey or larval food in Malaysia, Brazil, and Australia suggests these taxa are closely associated with stingless bees and may contribute to honey properties. By reducing the genomic underrepresentation of evolutionarily divergent sister clades related to *Nicoliella* and *Acetilactobacillus*, our genome-resolved analyses reveal a globally distributed, phylogenetically cohesive Lactobacillaceae species complex dominating pot honey.

## Introduction

For centuries, indigenous groups have managed stingless bees of the tribe Meliponini to produce pot honey [1, 2], which is recognized for their antimicrobial properties [3, 4], therapeutic potential [5, 6], and traditional use in treating ailments [6, 7]. Growing interest in stingless bee products such as honey [3, 4, 8], and propolis [9], highlights the potential of beekeeping to promote family [10] and community economic development [11, 12]. Beyond their economic value, stingless bees contribute significantly to ecosystem services, providing pollination of native crops [13, 14] in tropical regions. Meliponiculture, the keeping of stingless bees, supports local livelihood and safeguards biocultural heritage by integrating traditional ecological knowledge, biodiversity conservation, and cultural identity.

**Fig. 0.**
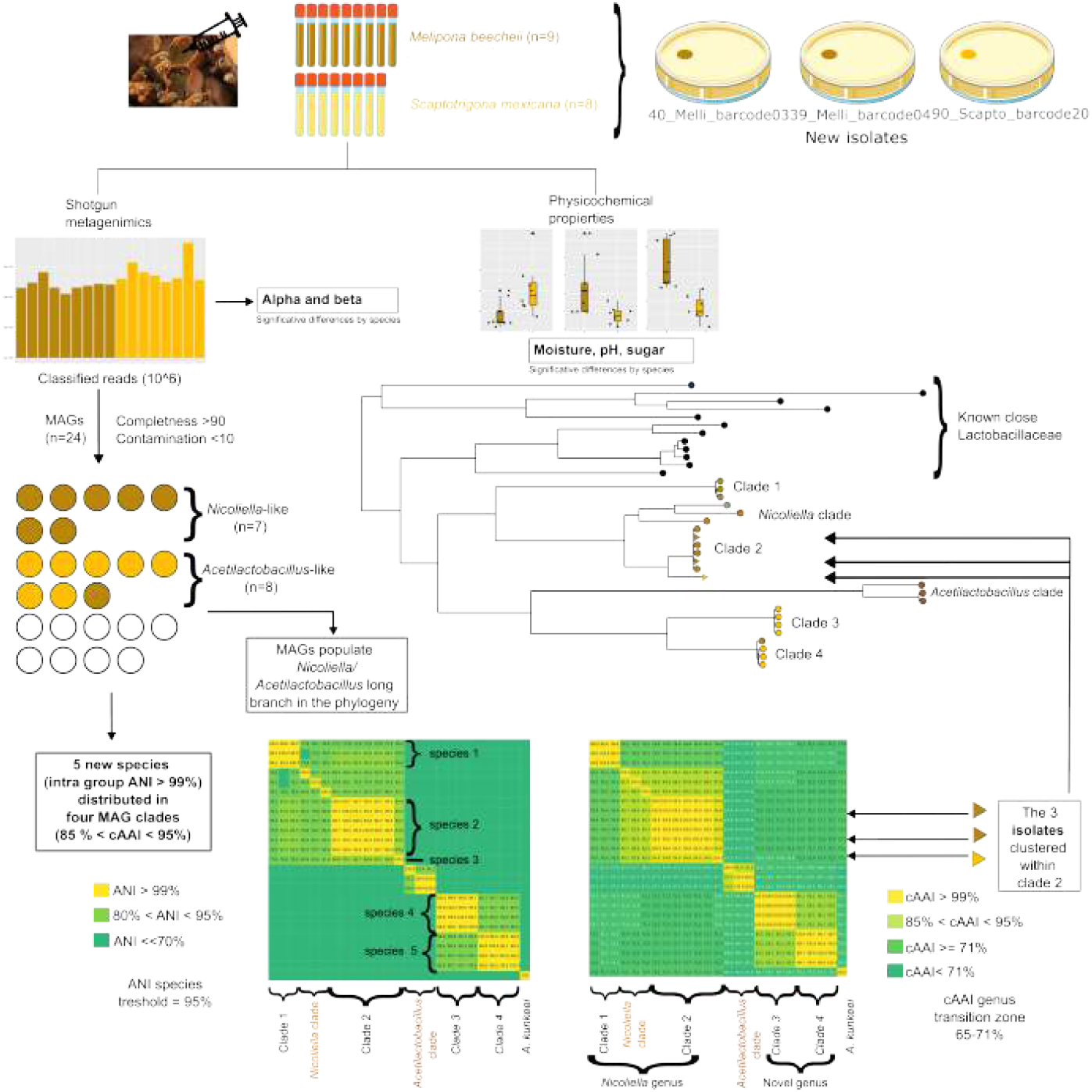
Genome-resolved analyses reveal that pot honey is dominated by a phylogenetically cohesive Lactobacillaceae species complex. From 17 samples, 15 MAGs and three isolates were clustered into four previously undescribed clades, forming five ANI-defined species and two cAAI-defined genus-level lineages related to *Nicoliella* and *Acetilactobacillus*. Physicochemical and microbiological properties separate seventeen honey samples.

The therapeutic effects of honey [15], and particularly of pot honey [5] have been attributed to its diverse bioactive compounds [16]. However, organisms isolated from pot honey also exhibit antimicrobial properties [17, 18, 19], making their identification particularly relevant. Metagenomics and metabarcoding have transformed bacterial ecology by enabling the characterization of microbial communities and the discovery of uncultivated species [20, 21]. In food-associated microbiomes shotgun metagenomics enables the reconstruction of metagenome assembled genomes (MAGs) providing insights into community composition [22, 23] and metabolic potential [24]. Although, the honey microbiome has recently gained attention, shotgun metagenomic studies have so far focused exclusively on *Apis mellifera* [*25, 26, 27, 28]*. *In contrast, the microbiota of stingless bees and their honey (pot honey) has been examined only trough microbial isolation [29, 30] and 16S metabarcoding [31]. Despite the advantages of shotgun metagenomics to asses genomic variability [32], and trace honey origin and authenticity [28], no such studies have been conducted on pot honey, leaving its microbial diversity largely unexplored*.

*Members of the Lactobacillaceae family are common components of both A. mellifera* [33] and Meliponini [34] bee microbiome, which has often been associated with bee health [35, 36, 30]. Lactobacillaceae also colonize honey, either produced by Meliponini [37] or by *A. mellifera* [25, 26, 28, 38]. Several Lactobacillaceae bacterial species have been identified in the honey and honey-crop through isolate-based 16S sequencing [30]. In pot honey, isolation [**?** 19] and a few metabarcoding studies [31] have shown the presence of two sister Lactobacillaceae genera: *Nicoliella* [**? ?**] and *Acetilactobacillus* [31]. *Nicoliella spurreliana* was isolated from the honey and homogenate of Australian stingless bee *Tetragonula carbonaria* [**?**], the other species in the genus is *Nicoliella lavandulae* isolated from lavandula [**?**]. *Acetilactobacillus jinshanensis* is the only known species [39] of its genus and was first isolated and consistently found in solid vinegar [40, 23, 41] having a recognized role in fermentation [42]. The identification of reported close relatives to *A. jinshanensis* such as *Lactobacillus sp*. Sy-1 isolated from Malaysian pot honey [43]—and the reconstruction of a MAG corresponding to *A. jinshanensis brasiliensis* in stingless bee larval food [44], suggest that the presence of a closely related *Acetilactobacillus*-like lineages ubiquitous to stingless bee-associated environments. However, assessing the microbial composition and its abundance distribution of pot honey remains largely unexplored.

To address this knowledge gap and to characterize the bacterial diversity we applied shotgun metagenomics in honey produced by two stingless Mexican bee species: *Melipona beecheii* and *Scaptotrigona mexicana* (Fig. 1). In seventeen pot honey samples our analyses revealed abundant Lactobacillaceae distributed in four clades with intraclade fastANI *>*99%. Our reconstructed MAGs share a maximum of 81% ANI with known public genomes, indicating a previously uncharacterized diversity of *Acetilactobacillus*-like species in these honey. Moreover, three strains isolated and sequenced from the same samples showed ANIs ranging from 84% to 99% to our MAGs confirming the presence of these previously uncharacterized microorganisms.

**Fig. 1.**
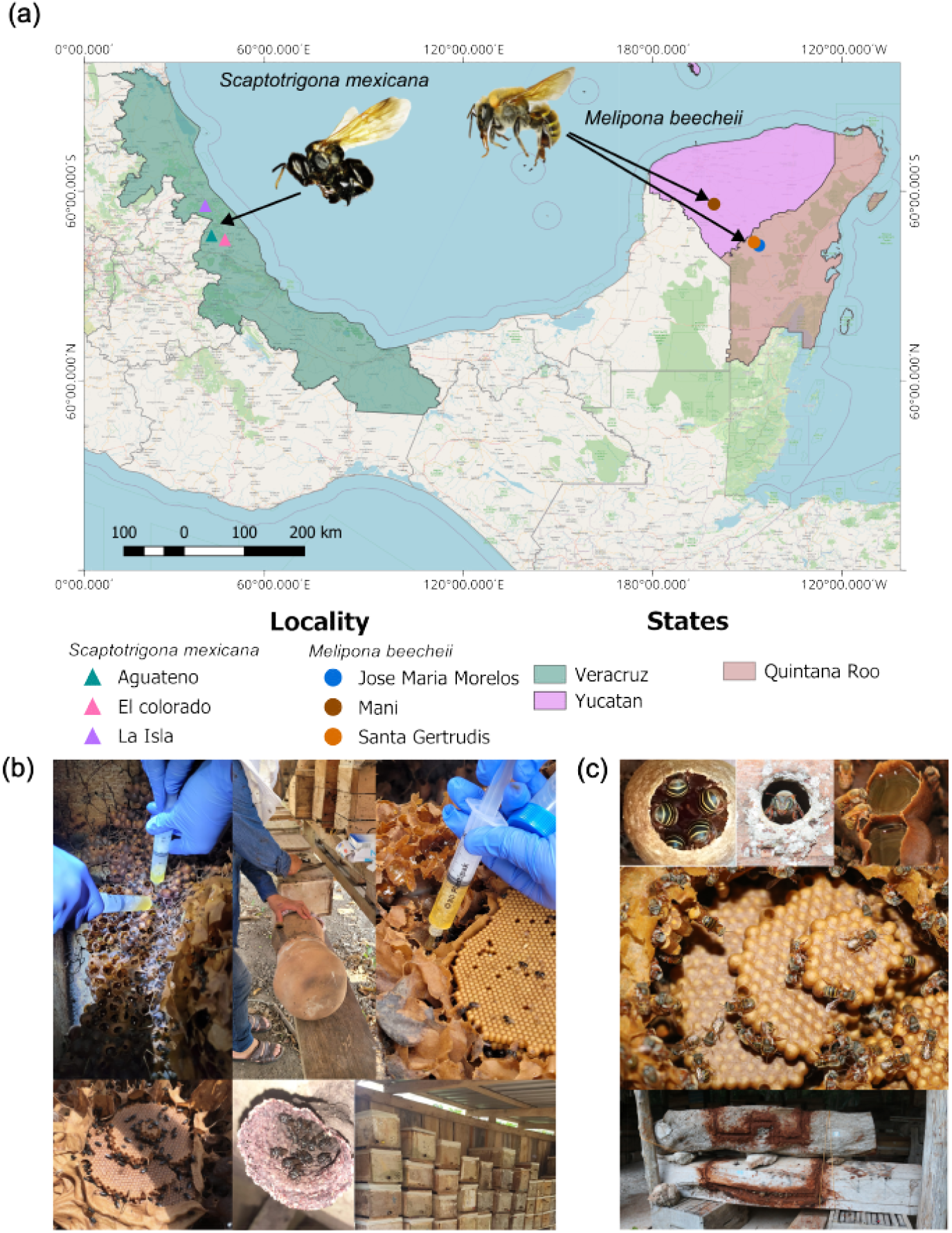
Honey from *S. mexicana* and *M. beecheii* were collected in Mexico. (a) Samples come from three states and six different localities. (b) Sampling *S. mexicana* honey from different zootechnical managements, including boxes and pitcher hives. (c) *M. beecheii* bees’ honey samples come from blended honey, beehive boxes, and traditional “jobones” beehive boxes, and traditional “jobones”, nest sections carved or cut from tree trunks, used historically as hives for stingless bees.

## Data summary

Metagenomes and genomes available in SRA BioProject PRJNA1235305. MAGs deposited on Zenodo:https://zenodo.org/records/18446951. Phylogenetic analysis available in https://microreact.org/project/8oNqsaGQm5fhFNgAiNobAe-honey-tree-v2. Code available at: https://github.com/HaydeePeruyero/honey_diversity. Authors confirm that all supporting data, code, and protocols have been provided in the article or through supplementary data files.

## Methods

### FIELD SAMPLING AND SAMPLE PROCESSING

#### Honey samples

During the 2022 harvest we selected meliponaries exploiting *M. beecheii* in the Yucatán Peninsula and *S. mexicana* in Veracruz (Table S1, Fig. 1a). Samples comprised several beekeeping methods: *S. mexicana* honey was harvested using modern boxes, blended honey, and traditional pitchers (Fig. 1b); *M. beecheii* using boxes, blends, and log hives (Fig. 1c). Meliponaries relied exclusively on their own resources during the previous year, with no supplementation of sugar or *A. mellifera* honey. Honey was aseptically extracted directly from closed pots using sterile syringes under strict hygiene conditions. For each sample, honey from one to three randomly selected colonies was pooled to a final volume of 320mL, divided into eight 40mL aliquots in sterile 50mL tubes, transported on ice at 10 − 15^°^C, and stored at −40°C at the CIDAS-QROO-UIMQROO. All samples were used in subsequent analyses.

#### Physicochemical Analyses

Parameters including moisture, pH, total sugars, electrical conductivity, ash content, color (Pfund scale), and hydroxymethylfurfural (HMF) were measured as described by Xolalpa [45], methodological details are provided in the supplementary material (Table S2).

#### Statistical analyses

were performed using a significance level of *α*=0.05 on 17 samples (9 *S. mexicana*, 8 *M. beecheii*). Normality (Shapiro–Wilk test and residual inspection) and homogeneity of variances (Levene’s and Bartlett’s tests) were assessed prior to each analysis. When assumptions were met, one-way ANOVA with Tukey’s HSD was used; otherwise, Kruskal–Wallis followed by Dunn–Bonferroni and Wilcoxon–Holm pairwise tests were applied, (Table S3). All analyses were conducted in R.

### DNA EXTRACTION AND PURIFICATION

#### Metagenomic DNA extraction

For each sample, 40g were pretreated with TissueLyser, following standard honey extraction protocols [46] [47]; Extensive details are provided in the supplementary material (Table S4). Subsequently, extraction was performed using the DNeasy Blood & Tissue^™^ Kit (Qiagen), following the manufacturer’s instructions. DNA yields exceeded 500ng, with concentrations >20ng/*µ*L and 260/280 ratios >1.7, indicating high purity (QIAxpert, Qiagen).

#### Shotgun Metagenomic Sequencing

DNA from seventeen honey samples (Table S1) was dried using an Eppendorf^®^ Vacuum Concentrator Plus (1hr) to remove residual water. Final concentrations ranged from 500–1000ng. Shotgun metagenomic sequencing was conducted at Macrogen, Inc. Libraries were prepared with the TruSeq Nano DNA Kit (Illumina) using 350bp inserts and sequenced on the NovaSeq 6000 platform, yielding 150bp paired-end reads.

#### Microbial isolation and extraction

Bacterial were isolated from honey samples following a protocol adapted from *Lactobacillus sp*. Sy-1 isolation [43] (Table S5). Isolates were preserved in MRS broth containing 20% (v/v) glycerol at −80°C. For biomass production, frozen stocks were revived on MRS agar and a single colony was inoculated into liquid MRS broth and incubated at 30°C for 24hr prior to cell harvesting and genomic DNA extraction. Genomic DNA was extracted following a phenol–chloroform protocol with minor modifications (Table S5).

#### Sequencing

For long-read sequencing, libraries were prepared using the Rapid Sequencing Kit V14, SQK-RAD114 Oxford Nanopore Technologies (ONT) according to the manufacturer’s instructions. Sequencing was carried out on a MinION device using a 24h sequencing run with default parameters, and data acquisition was performed using the MinKNOW graphical user interface (version 5.3.6).

### BIOINFORMATIC ANALYSES

#### Metagenomes

From quality control to diversity analysis, we followed “The Carpentries Metagenomics Pipeline”[48] modifying the assembly tool. Every metagenome have >6*×*10^7^ raw reads (Table S1). FastQC v0.1 [49] evaluated rawreads quality. Trimmomatic v0.39[50] `SLIDINGWINDOW:25:28, MINLEN:35, ILLUMINACLIP: TruSeq LT CD:2:40:15` removed low-quality nucleotides and adapters. Trimmed reads were assembled with MEGAHIT v1.2.9[51] (30 threads). For taxonomy classification, we used kraken2 v2.1.2 [52], and database PlusPFP (https://benlangmead.github.io/aws-indexes/k2) [53]. Results integrated in a biom file with Kraken-biom v.1.2.0[54]. **Diversity**. Beta diversity (Bray-Curtis distance) was visualized via NMDS, revealing between-sample variation. Alpha diversity, reflecting within-sample richness, was evaluated with Observed richness, Shannon, and Chao1 indices. Shannon accounts for abundance distribution, while Chao1 estimates unseen diversity from rare OTUs. Plots were generated in R v4.4.2 using phyloseq v1.50.0[55].

#### MAGs

We obtained MAGs with MaxBin v2.2 [56] and assessed their quality with CheckM v1.2.1[57]. All MAGs were refined using GC/coverage profiling (seqkit v2.11.0[**?**]; and then, clade-specific reference-guided mapping (minimap2 v2.30-r128[**?**]; bowtie2 v2.5.[**?**], BAM processing (samtools v1.18[**?**]) was performed for each clade, and the highest-quality, unrefined MAG served as the reference genome. Finally we applied iterative outlier-based filtering. Only refined MAGs with completness > 90% and contamination *<* 10% were retained (*n* = 24) (Table 1).

**Table 1.** Genomic features of MAGs with completeness > 90% contamination *<* 10% and all isolates.

| Genome ID | Group | Completeness | Contamination | Size (Mbp) | ANI to Closest known genome | Closest genus in GTDB | Max AAI to known genomes (*Nicoliella* or *Acetilactobacillus*) | Max core-gene AAI(%, *Nicoliella*) | Core positions compared |
| --- | --- | --- | --- | --- | --- | --- | --- | --- | --- |
| 40_Melli.029* | 1 | 93.1 | 1.72 | 1.4 | 78.14 | *Nicoliella* | 68.75 | 76.49 | 38027 |
| 45_Melli.016* | 1 | 91.38 | 1.72 | 1.7 | 78.20 | *Nicoliella* | 69.16 | 76.45 | 38303 |
| 46_Melli.022* | 1 | 91.38 | 3.45 | 1.5 | 78.13 | *Nicoliella* | 69.19 | 76.45 | 38303 |
| 22_Melli.029* | 2 | 98.28 | 2.04 | 1.9 | 80.48 | *Nicoliella* | 80.13 | 89.29 | 38289 |
| 40_Melli.barcode03 | 2 | 97 | n.a. | 1.9 | 80.78 | *Nicoliella* | 80.19 | 89.25 | 38300 |
| 45_Melli.020* | 2 | 98.28 | 2.51 | 2.1 | 80.35 | *Nicoliella* | 80.18 | 89.27 | 38251 |
| 47_Melli.001* | 2 | 97.41 | 4.31 | 2.1 | 80.18 | *Nicoliella* | 80.18 | 89.27 | 38252 |
| 39_Melli.barcode04 | 2 | 96 | n.a. | 1.9 | 80.65 | *Nicoliella* | 80.18 | 89.24 | 38243 |
| 41_Melli.001* | 2 | 98.28 | 1.72 | 1.8 | 80.39 | *Nicoliella* | 80.39 | 89.30 | 38269 |
| 90_Scpto.barcode20 | 2 | 87 | n.a. | 1.5 | 81.37 | *Nicoliella* | 80.64 | 88.90 | 37902 |
| 51_Scpto.004* | 3 | 98.28 | 0 | 1.3 | < 77 | *Acetilactobacillus* | 66.10 | 71.12 | 38201 |
| 89_Scpto.002* | 3 | 98.28 | 1.72 | 1.2 | < 77 | *Acetilactobacillus* | 66.08 | 71.11 | 38201 |
| 90_Scpto.012* | 3 | 98.28 | 3.61 | 1.2 | < 77 | *Acetilactobacillus* | 66.08 | 71.11 | 38201 |
| 92_Scpto.009 | 3 | 96.55 | 7.68 | 1.1 | < 77 | *Acetilactobacillus* | 66.18 | 71.13 | 38151 |
| 42_Melli.004* | 4 | 93.1 | 1.72 | 1.2 | < 77 | *Acetilactobacillus* | 65.96 | 71.57 | 38254 |
| 23_Scpto.001* | 4 | 91.54 | 1.72 | 1.2 | < 77 | *Acetilactobacillus* | 65.82 | 71.60 | 38244 |
| 89_Scpto.001* | 4 | 93.26 | 1.72 | 1.2 | < 77 | *Acetilactobacillus* | 65.81 | 71.58 | 38247 |
| 92_Scpto.002* | 4 | 93.1 | 4.31 | 1.2 | < 77 | *Acetilactobacillus* | 65.89 | 71.59 | 38244 |
| 39_Melli.018* | - | 99.69 | 4.36 | 3.2 | n.a. | *Enterobacteriaceae* | - | - | - |
| 43_Scpto.014* | - | 99.84 | 2.3 | 3.6 | n.a. | *Rosenbergiella epipactidis* | - | - | - |
| 43_Scpto.022* | - | 98.28 | 3.45 | 1.9 | n.a. | *Acetobacter* | - | - | - |
| 46_Melli.006* | - | 98.28 | 2.04 | 1.6 | n.a. | *Fructobacillus* | - | - | - |
| 46_Melli.021* | - | 100 | 0 | 2.3 | n.a. | *Bombella* | - | - | - |
| 92_Scpto.020* | - | 91.68 | 4.89 | 5.0 | n.a. | *Pantoea ananatis* | - | - | - |
| 46_Melli.018 | - | 100 | 5.17 | 2.8 | n.a. | Lactobacillaceae | - | - | - |
| 92_Scpto.034 | - | 94.83 | 9.25 | 1.3 | n.a. | *Fructobacillus* | - | - | - |
| 22_Melli.031 | - | 99.69 | 3.03 | 2.9 | n.a. | *Enterobacteriaceae* | - | - | - |

\* Indicates a high quality MAGs with < 5% of contamination and > 90% of completeness.
Note. Completeness of isolates was estimated with BUSCO.
Note. ANI, AAI, and cAAI have as maximum match *Nicoliella*, and this are the values reported.
Note. Values reported as < 77 in ANI correspond to values below the reporting threshold of FastANI.

#### Genome Assembly

Whole-genome sequencing of bacterial isolates was performed to validate MAG reconstruction. For each isolate, ONT adapters were removed from raw reads using Porechop 0.2.4 [**?**]. NanoStat 1.6.0 [**?**] assessed read length and quality statistics, while Filtlong 0.2.1 [**?**] removed the worst 1% of the reads. A de novo assembly approach was applied on genomes with depth >10X. First, Minimap 2.27[**?**] identified read overlaps (ava-ont preset). Then, Miniasm 0.3 [**?**] assembled the overlaps and Minipolish 0.2.0 [**?**] polished them. Assembly contiguity was assessed with the stats.sh script from BBTools 39.20 [**?**] while completeness was evaluated with BUSCO 5.5.0 [**?**] using *Lactobacillales* ODB10 as reference.

#### Phylogenetic analysis

For each MAG and genome we identified its closest match in the Genome Taxonomy Database (GTDB) with GTDB-Tk v2.5.2[58]. MAGs and genomes closest to *Nicoliella* or *Acetilactobacillus* were considered in posterior analysis. The core genome was calculated using Anvi’o v7[59] for MAGs (n=15), genomes from isolates (n=3), and the following public genomes: *Acetilactobacillus* (n=3), *Nicoliella* (n=2), *Lactobacillus sp*. Sy-1, and the Lactobacillaceae genomes closest to *Acetilactobacillus* according to the 2020 family reorganization (n=11) [39]. *Lentilactobacillus* senioris DSM24302 was included here as an outgroup, in accordance with its position on the 2020 reorganization phylogeny (Table S6). A **phylogenetic tree** was reconstructed using 129 core single-copy protein families obtained by setting (Anvio [59]), `anvi-gen-phylogenomic-tree`, specifically `anvi-get-sequences-for-gene-clusters` with `1=min=max number of genes per genome`). Sequences were aligned using MUSCLE v3.8.15 [**?**]. Evolutionary model was selected (`-m MFP`) in IQ-TREE v2.1.4-beta [60] with LG+F+R4 identified as the best-fit model according to the BIC criterion. Ultrafast Bootstrap support was applied with 1000 replicates (`-bb 1000`), and branch support was further evaluated using approximate likelihood ratio test (`-alrt 1000`).

#### Identity indexes

For each pair of MAGs and genomes in the pangenome FastANI v1.34 [61] computed ANI. Average Aminoacid identity (AAI) was calculated pairwise by EzAAI [**?**], while conserved Average Aminoacid Identity (cAAI) was approximated by calculating the aminoacid identity over the 129 conserved single copy protein families previously described. Results were visualized in a heatmap using Python v3.9.19 packages Seaborn v0.13.2 and Matplotlib v3.9.0.

#### Metapangenome

A metapangenome uses competitive mapping to align metagenomic reads against a set of genomes [62]; [63]. All *Acetilactobacillus*-like MAGs (n=8, 1 from *M. beecheii* and 7 *S. mexicana*), all *Nicoliella*-like MAGs (n=7 from *M. beecheii*), our three isolates, available *Nicoliella* genomes (n=2), *Lactobacillus* sp. Sy-1, available *A. jinshanensis* genomes (n=3), and one *A. kunkeei*, were included to create an Anvi’o v.8 [59] genome database. Open reading frames (ORFs) were identified via Prodigal v2.6.3 [64]. Gene families were calculated from the genome database using `anvi-pan-genome -ncbi-blast --mcl-inflation=10.` Honey metagenomes were mapped to MAGs and genomes with bowtie2 [65] with default parameters (--sensitive). Functional annotation across genomes was performed using ORFs and by annotating ORFs using BLASTp searches against the NCBI COG 2024 database [**?**]. Subsequently, the program `anvi-display-functions` identified differentially occurring functions across all genomes based on the COG24 PATHWAY annotation. To reduce noise and focus on conserved functional signals, only functions present in at least three genomes were retained `(--min-occurrence 3).`

## Results

### Honey from *M. beecheii* and *S. Mexicana* exhibited distinct physicochemical properties and microbial communities

Samples were grouped by bee species to compare their physicochemical and microbiological profiles. Significant differences were found in: pH, moisture and sugar content, (Fig. 2a). *M. beecheii* honey had an average pH of 4.65 ± 0.76, while *S. mexicana* averaged 3.88 ± 0.23 (Kruskal-Wallis test, K-W p-value=0.01), both within the reported range of 3.5 − 5.5 for pot honey [66]. Moisture content was higher in *S. mexicana* 25.12 ± 2.5% than in *M. beecheii* 22.75 ± 1.25% (K-W p-value=0.01) while reducing sugar concentration was lower in *S. mexicana* 372.28 ± 47.65g/L compared to *M. beecheii* 537.58 ± 8.86g/L (Tukey’s test, p-value=0.0007) (Table S2). No significant differences were found in electrical conductivity, ash content, color, or hydroxymethylfurfural levels.

**Fig. 2.**
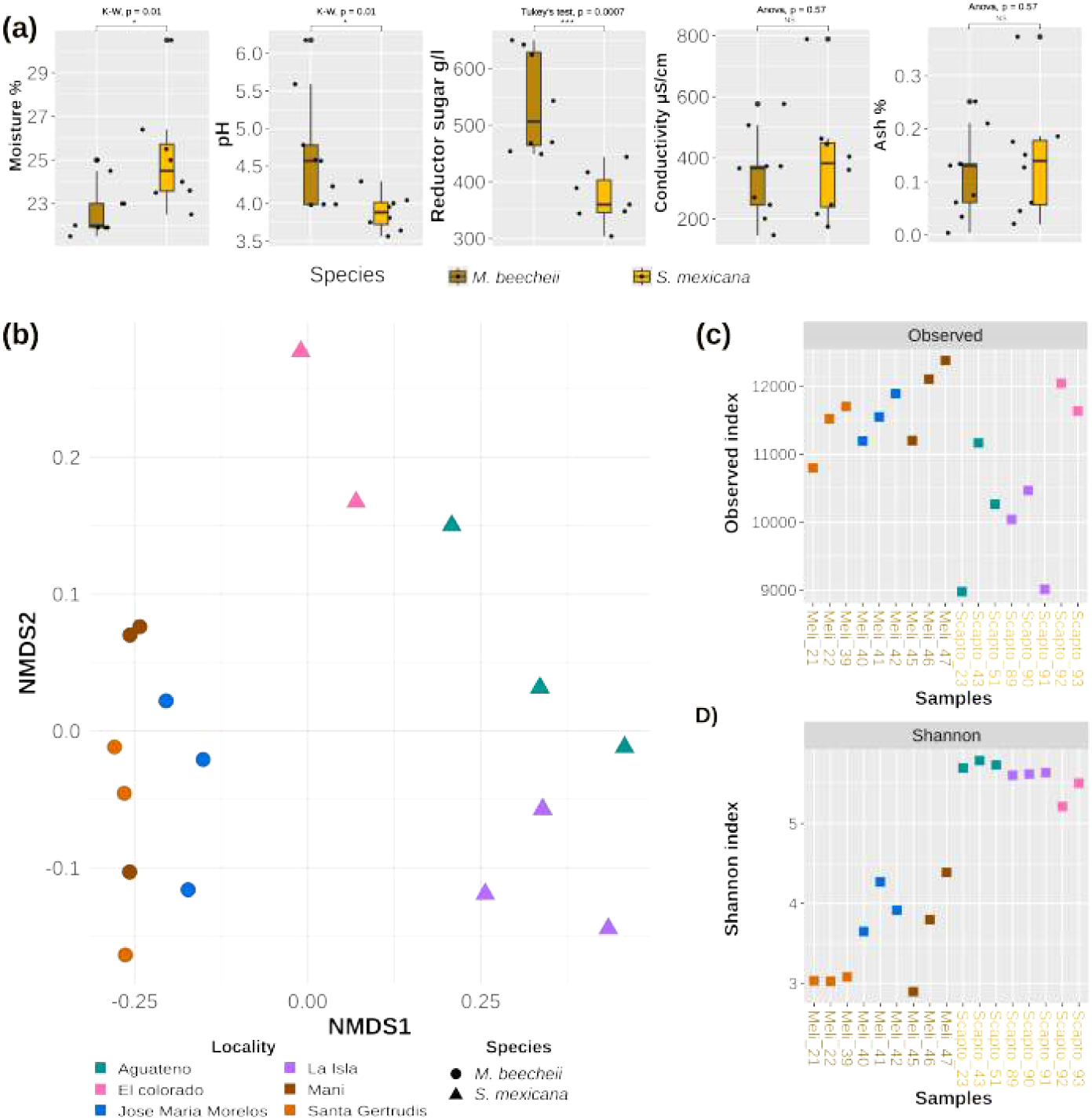
The diversity of honey’s microbial communities is grouped by bee origin. (a) Honey moisture, pH, sugars, conductivity, and ashes are shown in box plots according to species. (b) Species distribution and abundance are linearly separated in beta distance according to bee species. (c) *M. beecheii* has more alpha diversity than *S. mexicana*, measured by the observed index. (d) Contrary to taxonomic units, *S. mexicana* has more alpha diversity when measured by the Shannon Index.

Beta diversity separated samples by bee species (Fig. 2b), with tighter clustering in *M. beecheii* than in *S. mexicana*, (PERMANOVA pseudo-*F* = 42.362, p-value=0.0009; 1,000 permutations), indicating a strong species effect on honey microbiomes. In contrast, zootechnical practices did not affect clustering (Fig. S1a). Bee species and geography are confounded, as *M. beecheii* samples came from Yucatán and Quintana Roo and *S. mexicana* from Veracruz; thus, microbiological differences may also reflect variation in floral resources or honey physicochemical properties. Alpha diversity differed significantly between bee species with *M. beecheii* honey showing higher richness as measured by observed diversity (Welch t-test, p-value=0.025, Fig. 2c) and Chao1 (ANOVA with post hoc Tukey, p-value=0.0015, Fig. S1b) while *S. mexicana* showed higher evenness based on Shannon index (K–W, p-value=0.005, Fig. 2d). These patterns reflect differences in community evenness, with *S. mexicana* harboring more evenly distributed microbial communities and *M. beecheii* often being dominated by a single lineage.

### Previously undescribed *Nicoliella* /*Acetilactobacillus*-like Lactobacillaceae are consistently found in honey microbiome

Reads were classified using Kraken2, with ∼ 10^6^ reads assigned per sample (Fig. S2). Of these, 76.51% were bacterial, 20.52% eukaryotic, 2.82% viral, and 0.14% archaeal (Fig. 3a). Bacterial reads were dominated by Bacillota and Pseudomonadota (Fig. 3a). Fungi represented a minor fraction of the community, averaging 0.46% of reads (Fig. S4), and were mainly Ascomycota followed by Mucoromycota, Basidiomycota, and Microsporidia (Fig. 3b). Bacterial communities dominated the honey microbiome, with Lactobacillaceae as the most abundant family. Genera exceeding 5% relative abundance included *Acetilactobacillus, Apilactobacillus, Fructobacillus, Lactobacillus*, and *Nicoliella* (Bacillota), and *Pantoea* and *Tatumella* (Pseudomonadota) (Fig. 3c). Kraken-based taxonomic assignment indicated that *M. beecheii* honey was dominated by *Acetilactobacillus* (52.72%), whereas *S. mexicana* honey contained higher proportions of *Lactobacillus* (4.75%) and *Apilactobacillus* (9.77%) which has been found in flowers, bees, and hives [67].

**Fig. 3.**
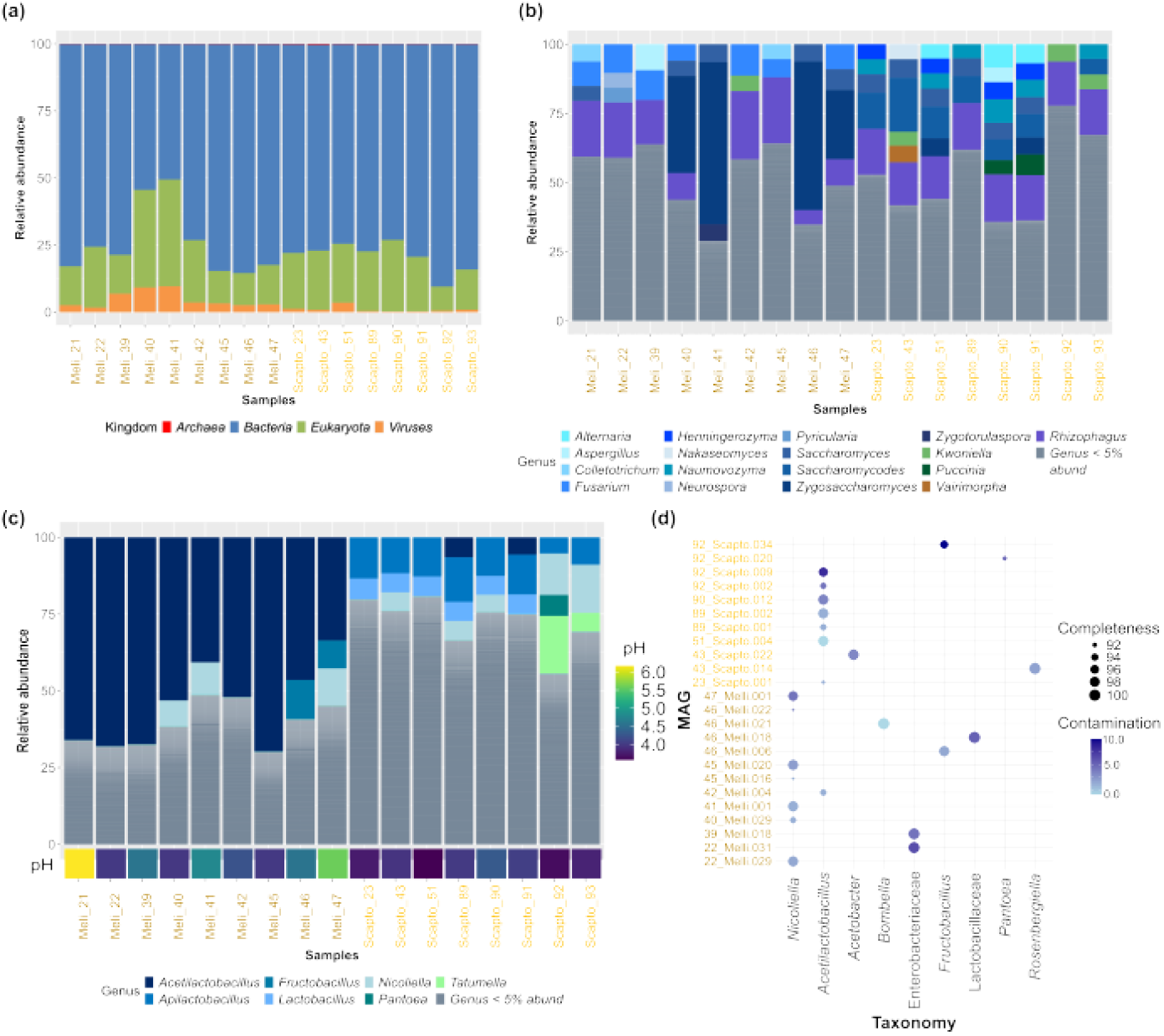
*Acetillactobacillus*-like lineage is consistently found in honey microbial communities. (a) Most classified reads are from the kingdoms of bacteria, followed by Eukaryota, and viruses. (b) Distribution of fungal genera, whereby those above 5% belonging to phyla Ascomycota are shown in blue, those belonging to *Basidiomycota* in green, those belonging to *Microsporidia* in brown, and those corresponding to *Mucoromycota* in purple. (c) Bacterial reads above 5% comprise the genera *Acetilactobacillus, Apilactobacillus, Fructobacillus, Lactobacillus*, and *Nicoliella*, from the family Lactobacillaceae within the Bacillota phylum (blue), and genera *Tatumella* and *Pantoea* within the *Pseudomonadota phylum* (green). (d) Twenty-four MAGs were recovered with more than 90% of completeness and less than 10% of contamination. Its closest genus, according to the GTDB, is on the x-axis, *Acetilactobacillus*

To improve species-level resolution, metagenomic bins were refined into MAGs and filtered to retain genomes with >90% completeness and *<*10% contamination, yielding 24 MAGs across both stingless bee species, of which 21 met MIMAG high-quality criteria (>90% completeness and *<*5% contamination) [**?**]. GTDB-based taxonomic assessment affiliated these MAGs with *Nicoliella, Acetilactobacillus, Rosenbergiella, Fructobacillus, Bombella, Pantoea*, and *Acetobacter* (Fig. 3d). In particular, the sister genera *Nicoliella* (n=7) and *Acetilactobacillus* (n=8) were assigned by GTDB as the closest relatives of fifteen of these MAGs; however, GTDB did not report ANI to any named species (Table1), reflecting the absence of sufficiently close reference genomes. Phylogenomic reconstruction and ANI clustering further resolved these genomes into four distinct, likely undescribed species-level clades (Fig. 5).

### Isolation of two *Nicoliella*-like lineages consistent with distinct ANI-based species

To provide experimental evidence for the presence in the honey of *Nicoliella*/*Acetilactobacillus*-like taxa, bacterial strains were isolated by picking individual colonies from dilution plates. To capture MAG diversity, colonies were isolated from samples yielding MAGs from each of the four clades: 40 Melli (clades 1–2), 90 Scapto (clade 3), and 89 Scapto (clade 4), as well as from sample 39 Melli, which yielded no MAGs. Isolates were sequenced using long-read Nanopore technology, and only datasets with >15× coverage were retained. Strains 40 Melli barcode03 and 39 Melli barcode04 clustered within MAG clade 2 (∼99% ANI), while 90 Scapto barcode20 showed its closest affiliation to clade 2 at lower similarity (85% ANI), (Table 1). These results demonstrate a close genomic correspondence between MAGs and cultured isolates, supporting the presence of two *Nicoliella*-like lineages consistent with distinct ANI-based species previously unclassified in the honey microbiome.

### Phylogeny resolves MAGs into four ANI-defined, previously uncharacterized species-level clades

To assess the phylogenetic placement of MAGs and isolates, we assembled a genome dataset comprising the closest relatives of the genera *Nicoliella* and *Acetilactobacillus*. We retrieved all *Acetilactobacillus* closest relatives (n=11) following the reclassification of Lactobacillaceae[39], we included as outgroup *Lentilactobacillus senioris* (Table S6). Public genomes included vinegar isolates *A. jinshanensis* HSLZ-75 [41] and *A. jinshanensis aerogenes*[68]; as well as *A. jinshanensis* brasiliensis, a MAG from stingless bee larval food [44]. We further added pot honey derived isolates related to *Acetilactobacillus*: *Lactobacillus* sp. *Sy-1* from Malaysian *Heterotrigona itama* honey [43], and *Nicoliella spurreliana* SGEP1 A5 [**?**] from Australian honey. In addition, we included *Nicoliella lavandulae* Es01 [**?**], a flower-derived isolate. Using this dataset, we reconstructed a multilocus phylogeny based on 129 core protein families (Fig. 4).

**Fig. 4.**
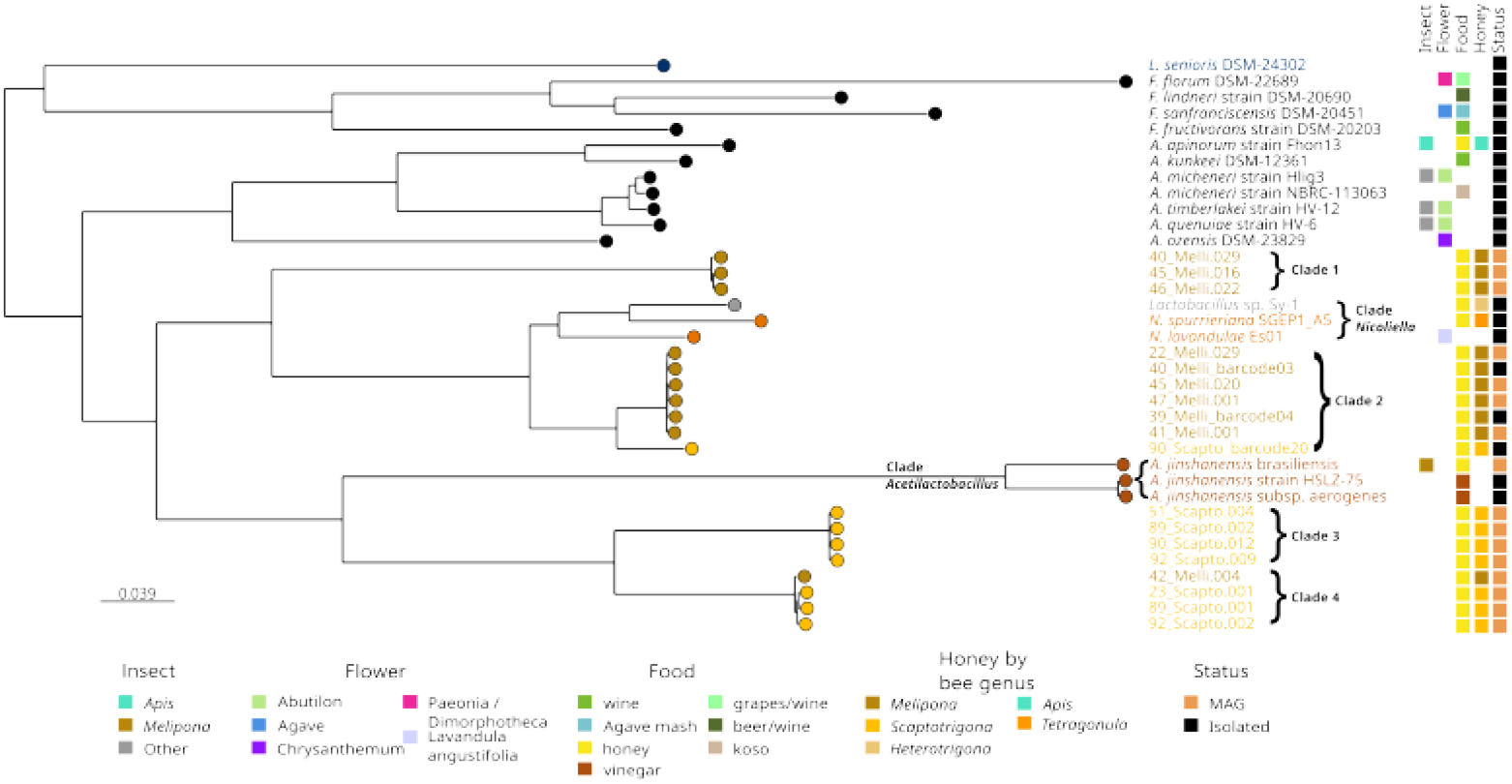
Phylogenomic tree based on 129 core proteins showing four clades of honey-associated Lactobacillaceae MAGs and isolates related to *Nicoliella* and *Acetilactobacillus*. MAGs cluster with pot honey–derived isolates from multiple geographic regions. Colors denote isolation source, honey-producing bee genera, and genome status (MAG or isolate). The phylogeny was reconstructed from an alignment of 40,955 amino acid positions using the LG+F+R4 substitution model. Branch support was assessed using ultrafast bootstrap with 1,000 replicates and the approximate likelihood ratio test

**Fig. 5.**
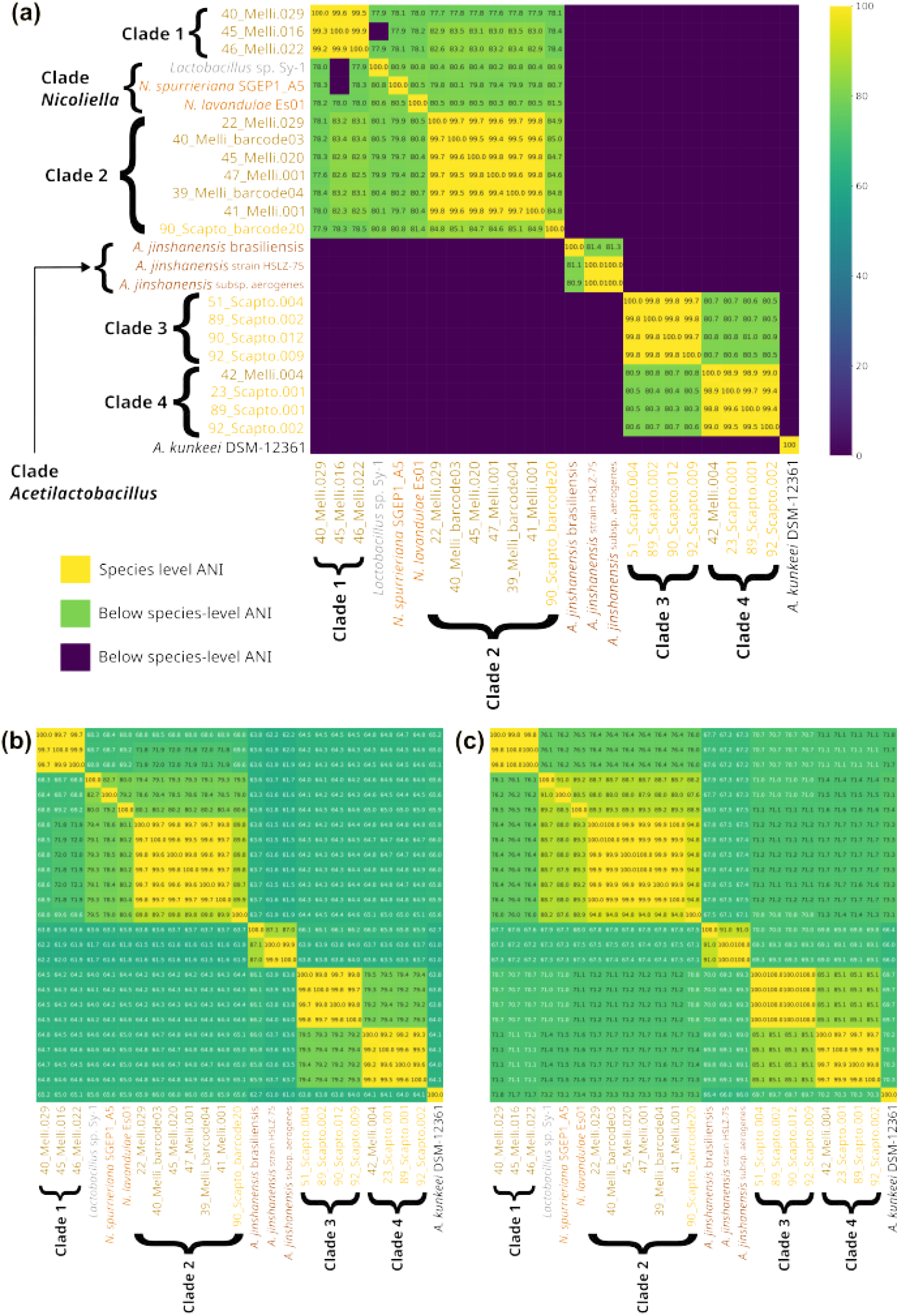
MAGs and isolates are clustered in four clades, spaning 5 species and 2 genus. **(a)** ANI values above 98% (highlighted in yellow) indicate species-level similarity, while interclade ANI values below ∼ 81% (green) reflect deep genomic divergence. MAGs and isolates cluster into four major clades, each showing high intraclade ANI (> 99%), except for Clade 2, where the inclusion of isolate 90 Scapto barcode20 results in lower intraclade ANI (∼ 96%). All MAGs exhibit ANI values < 81% relative to described Lactobacillaceae species, supporting their separation into multiple species-level lineages. **(b)** Clades 1, 3, and 4 display AAI values within the genus-level transition zone (≈ 65–71%) relative to *Nicoliella*, indicating substantial evolutionary divergence, whereas Clade 2 shows higher similarity. **(c)** Clades 3, and 4 fall below the genus-level transition range when compared to *Nicoliella* and *Acetilactobacillus* (65–71% cAAI). Together with phylogenomic and ecological evidence, cAAI supports Clades 3 and 4 as candidates for a previously uncharacterized genus.

Phylogenomic analysis grouped our MAGs and isolates into four clades, separating *Nicoliella* and *Acetilactobacillus* related genomes from other Lactobacillaceae members. *Nicoliella* genomes and *Lactobacillus* sp. *Sy-1* were closer to our MAGs Clade 2, while Clades 3 and 4 were closer to *Acetilactobacillus* clade, with all *Acetilactobacillus* forming a distinct lineage. A pair of genomes with ANI>95% are considered the same species;[39, 61], here, ANI values above 98% are shown in yellow (Fig. 5). All MAGs showed ANI<81% to any described Lactobacillaceae species, intraclade ANI exceeded 99% except in Clade 2 with 96% ANI due to isolate 90 Scapto barcode20 (Table S7), while interclade ANI values were <81% (Table S8). Excluding 90 Scapto barcode20 isolate from Clade 2 increased intraclade ANI to >99% and reduced variance to *≤*0.4%, resolving the genomes generated in this study into five species-level lineages, one represented solely by isolate 90 Scapto barcode20. Mean intra-clade ANI values for *Nicoliella* (87.11 ± 9.11%) and *Acetilactobacillus* (91.62 ± 9.34%) clades were below the 95% species threshold, indicating the presence of multiple species in both clades (Fig. 6). Among the three available *A. jinshanensis* genomes, the *brasiliensis* strain is the most phylogenetically divergent, with intra-clade ANI values below 81%, suggesting that its taxonomic placement may warrant reassessment.

**Fig. 6.**
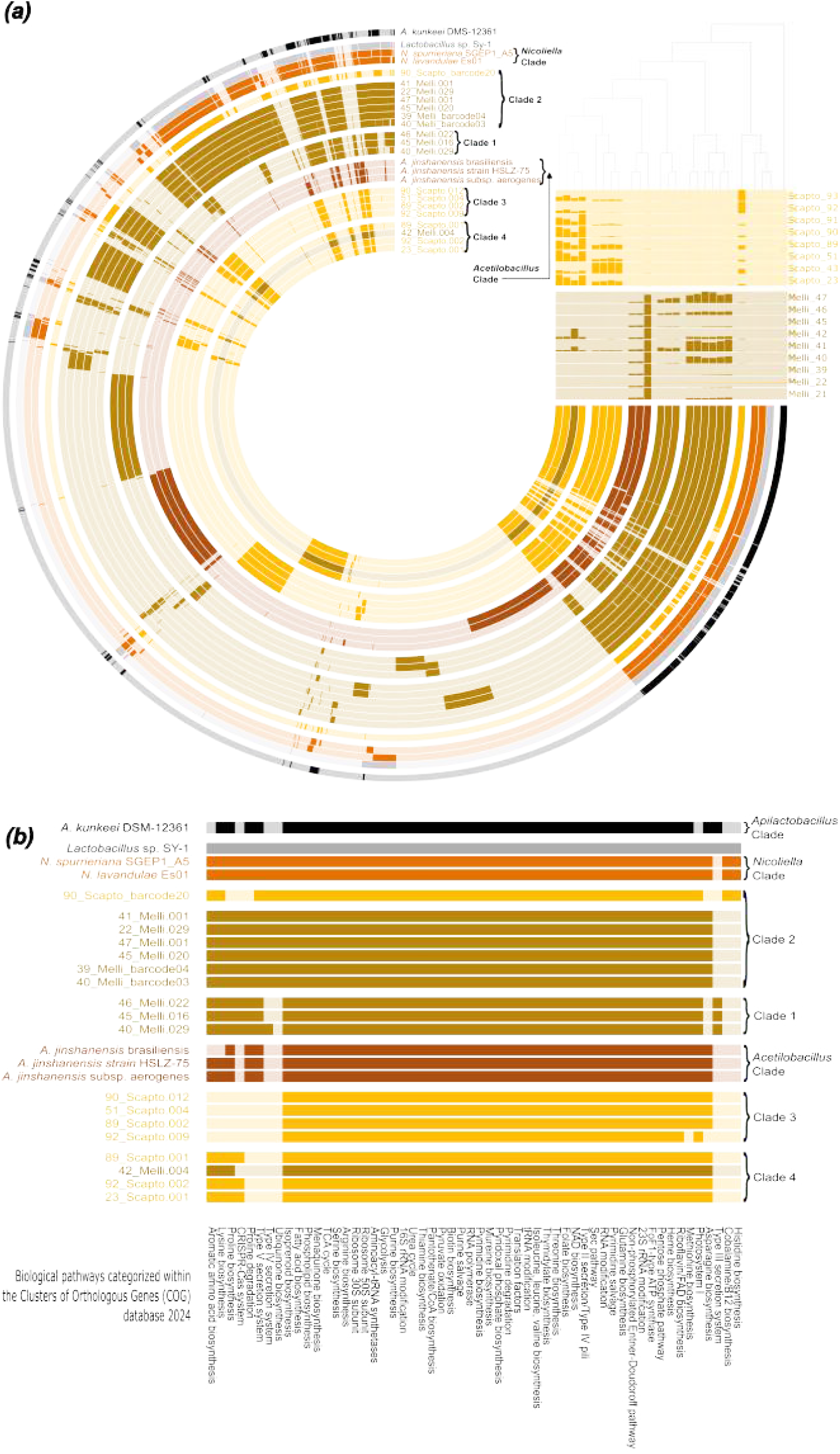
Metapangenome and functional analyses reveal clade-specific population structure and functions. (a) Presence–absence patterns of gene families cluster MAGs and isolates into four clades consistent with ANI and phylogeny, and distinct from *Nicoliella* and *Acetilactobacillus*. Competitive read recruitment highlights clade-specific mapping patterns across metagenomic samples. (b) Functional profiling based on COG categories shows clade-specific differences in amino acid biosynthesis and secretion systems.

### Genomic, ecological, and phylogenetic evidence for genus-level divergence

Criteria for defining novel genera in the Lactobacillaceae include: monophyletic clades, higher intra-than inter-genus AAI and cAAI with limited overlap, and shared ecological or physiological traits [39**?**]. All four MAG groups identified here form monophyletic clades in the phylogenomic tree. Clades 1 and 2 share a common ancestor with *Nicoliella*, whereas Clades 3 and 4 form a distinct lineage with a deeper common ancestor related to *Acetilactobacillus*. Clades 3 and 4 do not cluster within *Nicoliella* unless *Acetilactobacillus* is included, indicating substantial evolutionary divergence. Ecological and physiological differences further distinguish the clades. Isolates within clade 2 were obtained under culture conditions used for *Lactobacillus* sp. *Sy-1*, while no isolates were obtained within Clades 3 and 4. This suggests distinct physiological requirements consistent with clades 3 and 4 affinity to *Acetilactobacillus*. Genome size differences support clades separation, as Clades 3 and 4 have smaller genomes (∼1.2Mbp) than clades 1 (1.4–1.6Mbp), clade 2 (1.5–2.0Mbp), *Nicoliella* (∼2.0Mbp), and *Acetilactobacillus* (∼1.6Mbp) (Table 1).

In the Lactobacillaceae, inter- and intragenus cAAI values are typically separated by a transition zone of approximately 65–71% [**?**]. AAI values for Clades 1, 3, and 4 against their closest described genus, *Nicoliella* fall within this range, suggesting possible genus-level divergence. Inter clade values showed that Clade 1 (AAI= 68.83 ± 0.27%, cAAI= 76.31 ± 0.15%), Clade 3 (AAI= 64.35 ± 0.13%, cAAI= 71.06 ± 0.05%), and Clade 4 (AAI= 64.74 ± 0.26%, cAAI= 71.53 ± 0.05%) lie at the genus transition zone (Table S8, Fig. 5b). In contrast, Clade 2 displayed higher similarity to *Nicoliella* (AAI= 79.42± 0.86%, cAAI= 88.57 ± 0.66%) but remained clearly a distinct clade. Intraclade AAI and cAAI values were consistently high (≈97–99%), indicating strong internal coherence (Table S6, Fig. 5c).

Overall, Clades 1 and 2 are consistent with divergent species within a *Nicoliella*-related lineage, based on shared ancestry, high inter-clade cAAI (76–88%) against Nicoliella, and similar cultivation requirements. In contrast, Clades 3 and 4 show deeper phylogenetic divergence, lower AAI and cAAI values relative to *Nicoliella* (≈64%; ≈71% cAAI), smaller genomes, and distinct ecological signatures. Together, these features support Clades 3 and 4 as candidates for a previously uncharacterized genus within the Lactobacillaceae, pending formal taxonomic validation.

### Metapangenome reinforces ANI- and phylogeny-defined clades through population structure and functional differentiation

To evaluate the population structure of *Nicoliella*/ *Acetilactobacil* like MAGs and isolates in metagenomic reads, we constructed a metapangenome [62, 63] integrating read recruitment and gene family diversity (Fig. 6). The pangenome resolved core and variable protein families across genomes, and although the topology of the presence–absence-based cladogram differed from the phylogeny, MAGs and isolates largely clustered into the same clades, except for 90 Scapto barcode20, which grouped with *Nicoliella* genomes and *Lactobacillus sp*. Sy-1 instead of Clade 2. Reads from *M. beecheii* honey mapped mainly to MAGs in clades 1–2, with minimal recruitment to clade 4 and only one MAG in clade 3 (42 Melli), whereas *S. mexicana* honey mapped primarily to genomes in clade 4 and secondarily to the ones in clade 3. Among *A. jinshanensis* genomes, only *A. jinshanensis* brasiliensis recruited reads from *M. beecheii*, while other *A. jinshanensis* genomes showed little or no recruitment.

The main functional differences among clades involve amino acid biosynthesis and secretion systems. *Nicoliella* genomes encode complete pathways for the biosynthesis of histidine, asparagine, proline, lysine, and aromatic amino acids; in contrast, all MAGs and isolates lack histidine patwhay except for barcode20. Clade 1 additionally lacks asparagine patwhay, whereas Clade 3 and *A. jinshanensis brasiliensis* lack lysine and aromatic amino acid patwhay, with Clade 3 also mising proline patwhay. Ubiquinone degradation is absent in *Acetilactobacillus* and in Clades 1, 3, and 4. Regarding secretion systems, Clade 1 is the only group encoding a Type III, while Type V secretion systems are present in *Nicoliella, Acetilactobacillus*, and Clades 1 and 2; Clades 3 and 4 lack detectable secretion systems (Fig. 6).

## Discussion

In contrast to the well-studied honey microbiome of *Apis mellifera* [28] pot honey has been examined mainly through 16S ampicillins [31] and isolates [**?** 43] Using 17 shotgun metagenomes and three isolates, we showed that pot honey microbiomes are dominated by Lactobacillaceae [69] [25, 26, 28, 31]. Although Kraken2 classified most reads as *Acetilactobacillus*, genome-resolved analyses indicated that these reads represent previously uncharacterized *Acetilactobacillus*-like lineages.

Shotgun metagenomics yielded 14 high-quality MAGs (*≥*90% completeness and *≤*5% contamination) closely related to the sister genera *Nicoliella* (n=7) and *Acetilactobacillus* (n=7) but low ANI values to any named species (*≤*81%). Phylogenomic analysis based on 129 conserved protein families resolved MAGs and isolates into four monophyletic clades, two near *Nicoliella* and two near *Acetilactobacillus*. ANI-based clustering identified five species-level groups, with Clade 2 divided into two species: the first integrated by four MAGs and two isolates (intra clade ANI >99%) and the second represented by a single isolate, with ANI <84% to any genome. Together with Clade 1, these two species from Clade 2 represent three previously undescribed species-level groups closely related to the *Nicoliella* clade.

The 2020 revision of the former *Lactobacillus* lineage split it into 23 genera within the *Lactobacillaceae* [39]; *Nicoliella*, described in 2023 from pot honey, was therefore not included. In our dataset, the 3 species level groups in Clades 1 and 2 fall within the *Nicoliella* genus-level AAI/cAAI transition zone, consistent with possible affiliation with *Nicoliella* genus. In contrast, the 2020 revision included *Acetilactobacillus* as a genus represented by a single species, *A. jinshanensis*, characterized by a long evolutionary branch and sparse genomic representation; currently, only three genomes are available—two vinegar isolates and one MAG from stingless bee larval food. In our study, MAGs clades 3 and 4 clustered adjacent to the *Acetilactobacillus* clade. These *Acetilactobacillus*-like MAGs showed low AAI/cAAI values relative to available *A. jinshanens*is genomes supporting the interpretation that these MAGs represent a previously undescribed genus-level lineage and expanding the currently underrepresented *Acetilactobacillus*-related diversity. In the metapangenome reads from *M. beecheii* honey mapped preferentially to the *brasiliensis* genome and only marginally to other *A. jinshanensis* genomes, supporting the idea that honey harbors *Acetilactobacillus*-like lineages distinct from those associated with vinegar. This pattern suggests that previous reports of *Acetilactobacillus* in honey may reflect limitations of available reference genomes rather than true taxonomic identity.

Together, the recovery of three *Nicoliella*-related isolates and four closely related yet clearly distinct MAG clades distributed around *Nicoliella* and *Acetilactobacillus* clades, suggest that the apparent dominance of *Acetilactobacillus* in honey requires revision to account for previously unrecognized species within this lineage. Genome-resolved analyses further reveal that Lactobacillaceae dominance in pot honey reflects a complex population structure composed of multiple, closely related species-level lineages with broad geographic distribution.

Finally, ecological context may help explain the observed patterns. Honey represents a relatively acidic niche, consistent with the known physiology of *Acetilactobacillus*-related taxa. The inability to recover representatives of clades 3 and 4 as isolates may reflect more stringent growth requirements, such as higher acidity or anaerobic conditions, which is consistent with the significantly lower pH observed in honey from *S. mexicana* when compared to *M. becheii. A. jinshanensis* was originally isolated from solid-state vinegar, with an optimal pH range of 3.0–5.0 [41], and subsequent strains have been recovered from acidic fermented products, suggesting that related honey-associated lineages may require similarly conditions for cultivation.

Our *Nicoliella*-like MAGs and isolates cluster with honey- and stingless bee–associated isolates from Malaysia and Australia, as well as with a flower-derived isolate from Spain, whereas the *Acetilactobacillus*-like Clades 3 and 4 group with a MAG recovered from stingless bee larval food in Brazil. Functionally, these lineages differ in amino acid biosynthesis and secretion pathways, suggesting varying degrees of ecological specialization and host association. Future work focused on the formal taxonomic and physiological characterization should explore genomic signatures of specialized metabolism, including genes involved in production of antimicrobial compounds.

Taken together, our results indicate that the pot honey microbiome is dominated by a phylogenetically cohesive complex of closely related species with global distribution. From a human consumption perspective, the dominance and consistent presence of these lineages across geographically distant regions suggest a conserved microbial signature that could inform authenticity assessments, complementing physicochemical markers currently used to evaluate pot honey. The combination of phylogenetic coherence, ecological specialization, and broad geographic occurrence underscores their relevance in the honey environment and motivates further functional and taxonomic investigation.

## Conclusion

Pot honey is dominated by a group of phylogenetically cohesive Lactobacillaceae species complex with global distribution, uncovered through genome-resolved analyses.

## Author contributions statement

Conceptualization, conception, and design of study, writing, and submission of manuscript A.X.A. and E.J.D.S.; methodology. A.X.A., E.J.D.S., N.S.M and B.E.R.C.; analysis of DNA extraction and preparation for sequencing A.X.A, D.I.H.M, W.I.M.C; C.L.C. and L.R.O. isolate miscroorganisms; Statistical analysis, A.X.A. and H.C.P.; analysis of metagenomic data H.C.P., N.S.M and A.X.A; metapangenome analysis J.A.L.F; physicochemical analyses. A.X.A. and H.C.P.; writing and preparation of the original draft, A.X.A, N.S.M, H.C.P, and E.J.D.S.; writing and review and editing A.X.A, N.S.M, H.C.P, and E.J.D.S.; Writing of the article with contributions from D.I.H.M., W.I.M.C., B.E.R.C, J.F.R.C., F.B.G., B.E.O.V., H.C.P.; Management and acquisition of financial funds for the project, A.X.A, and E.J.D.S. All authors have read and accepted the published version of the manuscript.

## Acknowledgments

Our gratitude to the beekeepers for providing honey samples from their valuable hives and to Emil-Neme and Edith-Pazos for their sampling support. We thank Carlos-Ucan, Omar-Sánchez, Araceli-López, Rubén-Busambra, Cristian-Suárez, and Areli-Chimal from CIDAS-QROO/UIMQROO for their assistance in fieldwork, physicochemical analyses, and DNA extraction, Jorge-Cime and Grecia-Hernández for the technical support at CISEI/INSP. Thanks to Ángel-Chi for his assistance in analyzing territorial data, to Ricardo-Ayala, Ruben-Busambra, Cristian-Suárez, Areli-Chimal, Juan-Mayo, and Omar-Sánchez for their bee photographs to Anton-Pashkov for his bioinformatic support.

