## supplementary for "Genome-resolved metagenomics reveals a phylogenetically cohesive *Acetilactobacillus*-like species complex dominating stingless bee pot honey"

**Supplementary material of Uncharacterized *Acetilactobacillus*-like genera represent the most abundant microorganisms in honeys produced by native stingless bee species, *Melipona beecheii* and *Scaptotrigona mexicana***

### 1. Honey samples

Table S1. Origin of honey samples, species, state and locality, zootechnical management used in native communities in Mexico. Total read content considers both forward and reverse raw reads.

| *Species bee* | *Sample Id* | *Zootechnical Management* | *State* | *Locality* | *GPS_X* | *GPS_Y* | *Read count (10^7)* | *GC %* |
|---|---|---|---|---|---|---|---|---|
| *M. beecheii* (n=9) | Melli_21 | Blended honey | Quintana Roo | Santa Gertrudis | 19.80 | -88.77 | 6.632 | 42.0 |
| | Melli_22 | Log hive (Jobon) | Quintana Roo | Santa Gertrudis | 19.80 | -88.77 | 7.726 | 41.1 |
| | Melli_39 | Beehive box | Quintana Roo | Santa Gertrudis | 19.80 | -88.77 | 7.758 | 42.4 |
| | Melli_40 | Beehive box | Quintana Roo | Jose Maria Morelos | 19,75 | -88.70 | 7.372 | 39.8 |
| | Melli_41 | Blended honey | Quintana Roo | Jose Maria Morelos | 19.75 | -88.70 | 6.655 | 38.9 |
| | Melli_42 | Log hive (Jobon) | Quintana Roo | Jose Maria Morelos | 19.75 | -88.70 | 7.043 | 40.5 |
| | Melli_45 | Beehive box | Yucatan | Mani | 20.39 | -89.39 | 7.138 | 40.8 |
| | Melli_46 | Log hive (Jobon) | Yucatan | Mani | 20.39 | -89.39 | 6.806 | 41.6 |
| | Melli_47 | Blended honey | Yucatan | Mani | 20.39 | -89.39 | 6.961 | 40.5 |
| *S. mexicana* (n=8) | Scapto_43 | Traditional Pitcher | Veracruz | Aguateno | 19..90 | -97..14 | 7.538 | 33.3 |
| | Scapto_23 | Blended honey | Veracruz | Aguateno | 19.90 | -97.14 | 7.009 | 33.2 |
| | Scapto_51 | Beehive box | Veracruz | Aguateno | 19..90 | -97..14 | 7.729 | 34.1 |
| | Scapto_89 | Traditional Pitcher | Veracruz | La Isla | 20.36 | -97.24 | 7.149 | 33.9 |
| | Scapto_91 | Beehive box | Veracruz | La Isla | 20.36 | -97.24 | 7.503 | 33.9 |
| | Scapto_90 | Blended honey | Veracruz | La Isla | 20.36 | -97.24 | 7.381 | 34.4 |
| | Scapto_92 | Beehive box | Veracruz | El colorado | 19.84 | -96.94 | 7.739 | 40.2 |
| | Scapto_93 | Blended honey | Veracruz | El colorado | 19.84 | -96.94 | 6.628 | 37.4 |

### 2. Physicochemical parameters of honey samples

Honey samples were analyzed in accordance with standard procedures established by the Codex Alimentarius and AOAC methods. Parameters were measured as described by (Xolalpa *et al.,* 2024) as follows: **Moisture content** was determined using a digital RHB model 32ATC refractometer, with results with temperature-corrected refractive-index values. **pH** measurements were performed directly using an Okthon® Meter potentiometer equipped with a pH/ATC electrode (WD-35811-71) after diluting the honey samples with distilled water. **Reducing sugar content**. This was determined using a reflectometer (Merck RQflex 10; Darmstadt, Germany), which measures the combined glucose and fructose content using Reflectoquant reactive test strips (Merck Millipore™; Darmstadt, Germany). **Electrical conductivity** was measured with an Okthon® Meter conductivity meter. Electrical conductivity was determined at 20 °C using a conductivity meter following the methodology described by (Piazza *et al.*, 1991). The Okthon® potentiometer was used to calculate the **ash percentage** according to the formula C = (0.14 + 1.71) / A (Piazza et al.,1991). **Color** was measured using a Hanna HI96785 honey color photometer and expressed in Pfund units (mm). **Hydroxymethylfurfural (HMF)** was measured using the RQFlex 2.0 reflectometer with HMF test strips, according to the manufacturer's procedure. All analyses were performed in duplicate to ensure accuracy and reproducibility.

Table S2. Evaluation of the physicochemical parameters and their relationship with zootechnical management in honey from three species of bees.

| Species | State | Zootechnical management | Sample number (n) | Moisture % | Color (mm Pfund) | pH | Electrical conductivity (µS/cm) | Ash % | Reductor sugar g/L | HMF (mg/L) |
|---|---|---|---|---|---|---|---|---|---|---|
| | | | | Means | | | | | | |
| ***M. beecheii* (n=9)** | Quintana Roo | Beehive box | 2 | 24.0 | 53.5 | 4.28 | 196.45 | 0.03 | 592.50 | 2.85 |
| | | Jobon | 2 | 21.70 | 48.0 | 4.11 | 388.75 | 0.14 | 552.0 | 2.20 |
| | | Mixed Honey | 2 | 22.45 | 48.25 | 5.48 | 283.50 | 0.08 | 458.50 | 2.55 |
| | Yucatán | Beehive Box | 1 | 24.50 | 44.00 | 3.99 | 372.50 | 0.13 | 624.00 | 1.00 |
| | | Jobon | 1 | 22.00 | 43.00 | 4.57 | 576.50 | 0.25 | 470.00 | 1.00 |
| | | Mixed Honey | 1 | 21.90 | 43.00 | 5.59 | 372.50 | 0.13 | 538.50 | 1.00 |
| ***S. mexicana* (n=8)** | Veracruz | Beehive box | 3 | 26.37 | 33.83 | 3.81 | 274.93 | 0.08 | 355.00 | 1.00 |
| | | Cantaro | 2 | 24.00 | 63.00 | 3.98 | 453.75 | 0.18 | 374.50 | 4.55 |
| | | Mixed Honey | 3 | 24.63 | 47.17 | 3.89 | 455.33 | 0.18 | 388.43 | 0.97 |

Table S3. Average of physicochemical parameters of honey samples

| Specie | Moisture | pH | Reducing sugar | Electrical conductivity | Ash | Color | HMF |
|---|---|---|---|---|---|---|---|
| *M. beecheii* | 22.74 ±1.25 | 4.65±0.76 | 537.5±88.86 | 339.87±140.03 | 0.11±0.08 | 48.05±8.9 | 2.02±1.24 |
| *S. mexicana* | 25.12±2.50 | 3.88±0.23 | 372.28±47.65 | 387.28±194.83 | 0.14±0.11 | 46.12±16.3 | 1.87±2.20 |

High electrical conductivity can complicate DNA extraction, likely due to trace minerals in honey (0.17%), with potassium being the most abundant (Olaitan *et al.,* 2007). The electrical conductivity of *A. mellifera* honey from Thailand has been estimated at 0.26 ± 0.04 mS/cm (Wanjai *et al.,* 2012). In our study, *M. beecheii* honey had an average electrical conductivity of 339.87 ± 140.03 μS/cm, while *S. mexicana* honey averaged 387.28 ± 194.83 μS/cm (Table S2, S3), (ANOVA p-value = 0.56). The mean ash content was 0.11 ± 0.08% for *M. beecheii* and 0.14 ± 0.11% for S. mexicana (ANOVA, p-value = 0.56). In terms of colorimetry, the average values were 48.05 ± 8.9 for *M. beecheii* and 46.12 ± 16.35 for S. mexicana (K-W, p-value = 0.77). Lastly, HMF content averaged 2.02 ± 1.24 for *M. beecheii* and 1.87 ± 2.20 for *S. mexicana* (K-W, p-value = 0.47).

The range of hydroxymethylfurfural (HMF) for all samples was 1 to 4.55 mg/L (Table S2). This parameter is typically used to determine whether the honey has been heated. Souza *et al.,* (2021) report that Melipona scutellaris honey collected from different sites in northeastern Brazil exhibits variation in HMF content across samples. This geographic variation adds another layer of complexity to the study of HMF levels in honey, which has historically been considered in various studies of *A. mellifera* honey.

The Pfund scale is a standardized system for classifying honey color in *A. mellifer*a. Honeys showed Pfund readings of 33.8 mm in the "Beehive box", 63.0 "Cantaro", and 47.1 mm in the "Mixed honey" of *S. mexicana.* For *M. beecheii,* Pfund readings were 44.0 to 53.5 mm in "Beehive box," 43.0 to 48.0 mm in "jobon," and 47.1 to 48.25 mm in "Mixed honey " (Table S3). The range of honey colors for stingless bee honey is from 33.83 to 63 mm and an average of 47.32 mm with shades ranging from extra light amber and light amber, it should be taken into account that the color of honey is closely related to the botanical origin; Many factors can influence the color, some of them can be the environment, the season, the minerals, the Maillard reaction, the phenolic content, the pollen or the wax used, floral origin and storage duration can directly impact the coloration (Raweh *et al.,* 2023).

In this study, the ash content of the honey samples ranged from 0.03% to 0.25% (Table S2). Both species of native stingless bees studied here share their values, even considering the different zootechnical management. Ash percentages are linked to electrical conductivity, which, according to (Pauliuc *et al.,* 2020) and (Serrano *et al.,* 2004), helps determine their floral and geographical origins.

### 3. DNA extraction of honey

Table S4. Detailed pretreatment and extraction protocols of DNA from microbiomes

| | *Protocol* |
|---|---|
| *Pretreatment* | Protocols from (Soares *et al.,* 2015) and (Waiblinger *et al.,* 2012) consist on dispensing 40ml of honey from each sample into 4 sterile 50 mL Falcon tubes (12.5 grams of honey per tube). The tubes were centrifuged at 11,000 × g for 10 minutes. The supernatants were discarded, and the pellets were diluted in 5 mL of ultrapure water and pooled into a single Falcon tube (approximately 40 mL total). This reconcentrate was centrifuged again at 11,000 × g for 10 minutes. Again, the supernatant was discarded, and the pellet was resuspended in 0.5 mL of ultrapure water. This suspension was transferred to a 2 mL reaction tube containing tungsten beads (500 mm size). 200 µL of PBS was added to the suspension, which was then placed in the TissueLyser II (Qiagen) at 30 Hz for 4 minutes to achieve homogenization. |
| *Extraction* | **Extraction** was performed using the DNeasy Blood & Tissue™ Kit (Qiagen), following the manufacturer's instructions: For each honey sample, 40g was pretreated following this protocol: Extraction required Qiagen DNeasy Blood&Tissue. Lysis mix included 650µL mixture, 180µL ATL buffer, 20µL Proteinase-K, and 200µL ethanol, followed by purification using DNeasy spin columns. Elution was performed with 25 µL of AE buffer after centrifugation at 6000 g. The mixture was incubated for one minute, then centrifuged at 6000 g for one minute twice. DNA quantification was performed using a QIAxpert spectrophotometer (Qiagen). |

### 4. Bacterial Isolation and Sequencing

Table S5 Detailed pretreatment and extraction protocols of bacterial isolates

| | |
|---|---|
| isolation | The protocol was adapted from *Lactobacillus* sp. Sy-1(Syed Yaacob *et al.,* 2022), consisting of 1g of honey suspended in 10 mL of MRS broth, vortexed for 10s, and serially diluted ten-fold. Aliquots (100 µL) were spread onto MRS agar supplemented with 20 g/L fructose and 0.8% (w/v) $CaCO_3$, and the plates were incubated aerobically at 30°C for 3–4 days. Colonies producing clear zones indicative of $CaCO_3$ hydrolysis by lactic acid production were selected and subcultured to obtain pure isolates. |
| *Extraction of Genomic DNA* | Bacterial pellets were resuspended in 500 µL of lysis buffer containing 50 mM Tris-HCl (pH 8.0), 50 mM EDTA, and 3% SDS, which had been heated to 65 °C, and incubated at 65 °C for at least 40 min with occasional mixing. Subsequently, 500 µL of phenol:chloroform:isoamyl alcohol (25:24:1) was added, and the samples were mixed by inversion and centrifuged at 13,000 rpm for 5 min. The aqueous phase was transferred to a new tube, and 100 µL of 3 M sodium acetate was added, then gently mixed. DNA was precipitated by adding 1 mL of absolute ethanol and incubating the samples on ice for 25 min. After centrifugation at 13,000 rpm for 15 min, the supernatant was discarded, and the DNA pellet was washed with 400 µL of 70% ethanol. Samples were centrifuged at 13,000 rpm for 2 min, air-dried at 37 °C, and finally resuspended in 100 µL of water. DNA samples were stored at −20 °C until further use. |

### 5. Alpha and beta diversity

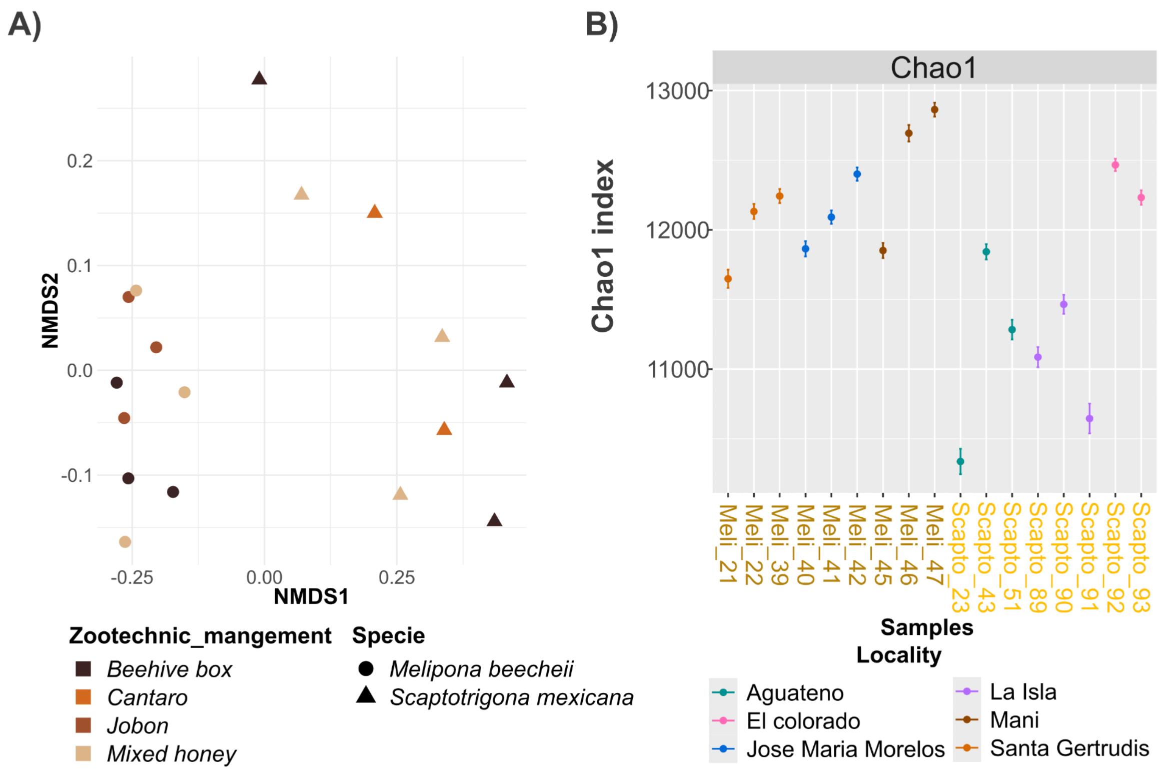

Fig S1 A) Beta diversity by zootechnic management. B) Chao1 diversity index of bacteria.

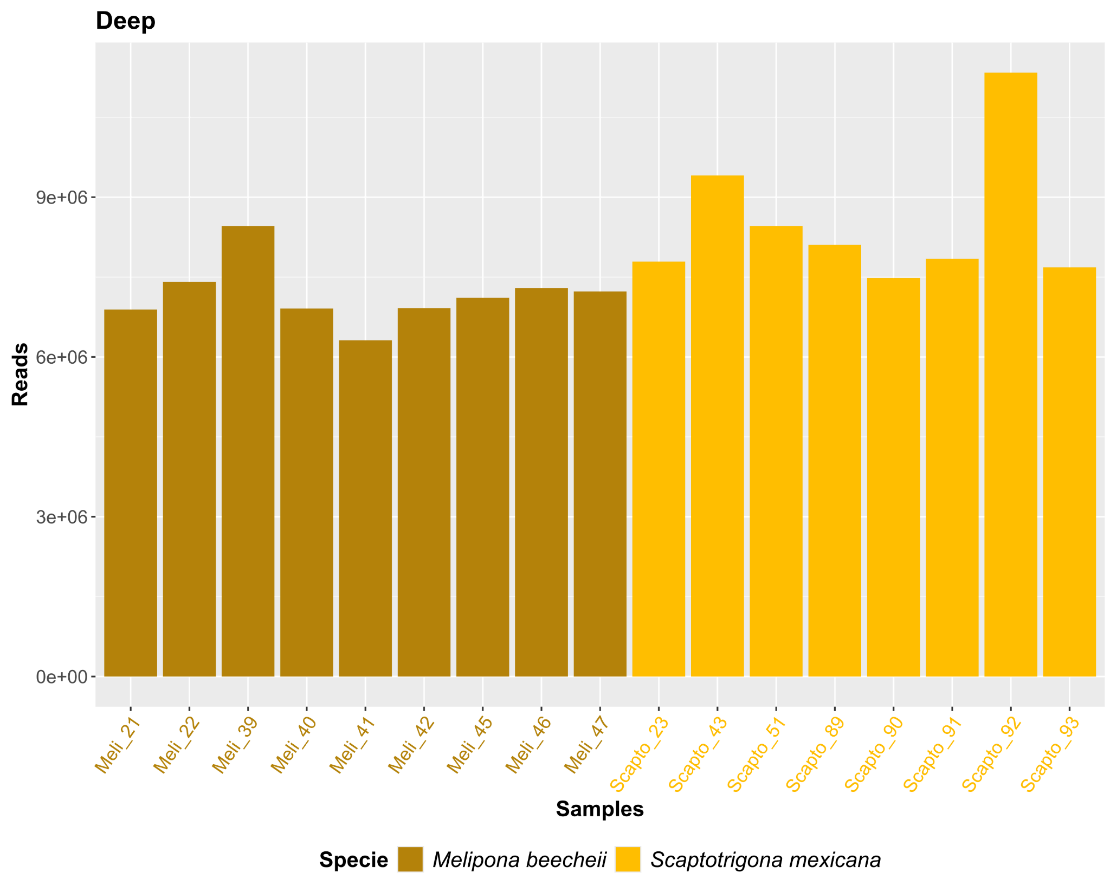

Fig S2. Samples have more than six million classified reads.

### 6. Taxonomical distribution

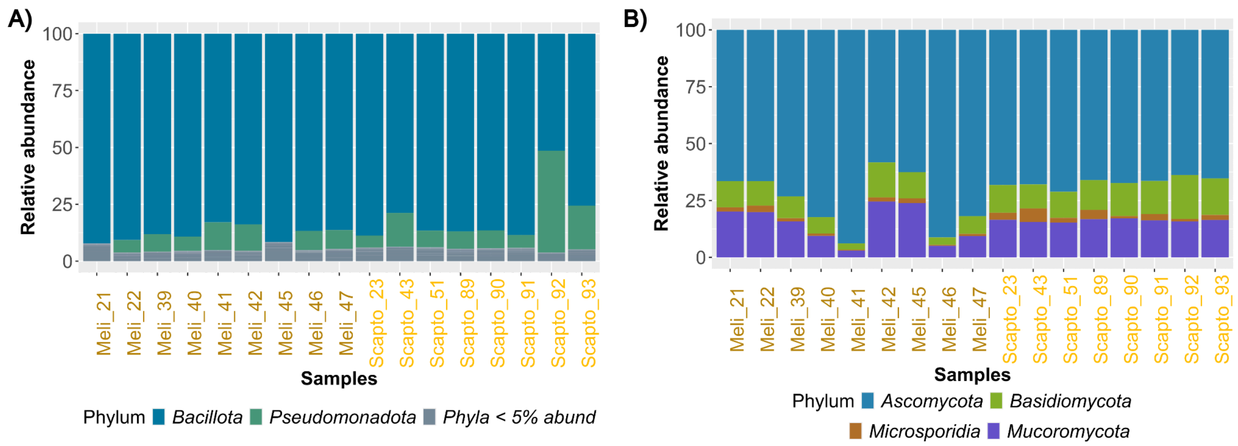

Fig S3 **A)** Bacillota is the most abundant bacterial phylum. Pseudomonadota is not highly represented in some M. beecheii samples. **B)** The two most abundant fungi are Ascomycota and Basidiomycota.

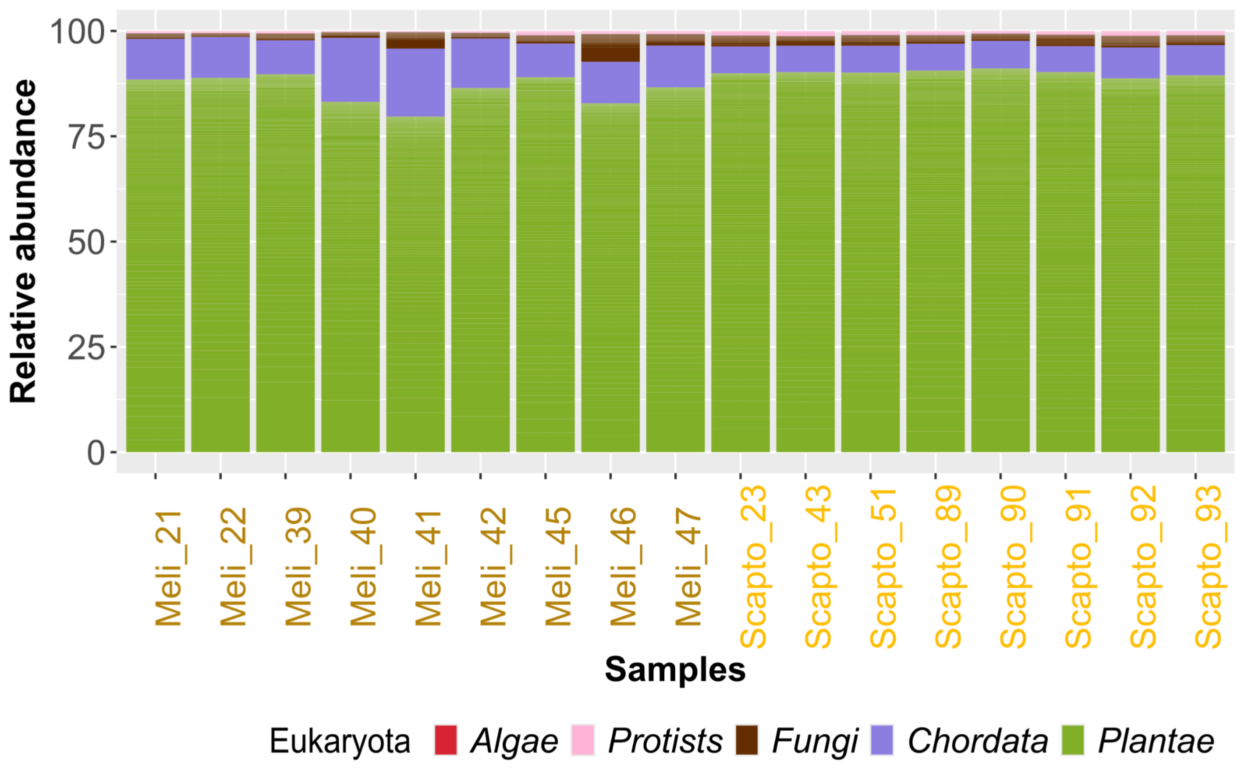

Fig S4. Within Eukaryota, the most abundant group is Plantae, followed by Chordata and Fungi. Fungi make up only 0.4% on average.

### *7. Potentially new species of Acetilactobacillus* are found in the honey microbiome.

Table S6. Public accessions of the 11 *Lactobacillus* in Zheng *et al*., 2020 classification that are closest to *Acetilactobacillus jinshanensis*. Additionally, Lactobacillus sp. Sy-1 is included.

| | **Genome sequence accession number:** | **16S rRNA gene accession number:** |
|---|---|---|
| *Lentilactobacillus senioris* | AYZR00000000 | AB602570 |
| *Apilactobacillus micheneri* | POSO00000000 | KT833121 |
| *Apilactobacillus kosoi* | BEXE01000000 | LC318484 |
| *Apilactobacillustimberlakei* | POST00000000 | KX656650 |
| *Apilactobacillus quenuiae* | POSN00000000 | KX656667 |
| *Apilactobacillus apinorum* | JXCT00000000 | JX099541 |
| *Apilactobacillus kunkeei* | AZCK00000000 | Y11374 |
| *Apilactobacillus ozensis* | AYYQ00000000 | AB572588 |
| *Acetilactobacillus jinshanensis* | CP034726 | KT783533 |
| *Fructilactobacillus sanfranciscensis* | AYYM00000000 | X76327 |
| *Fructilactobacillus lindneri* | JQBT00000000 | X95421 |
| *Fructilactobacillus florum* | AYZI00000000 | AB498045 |
| *Fructilactobacillus fructivorans* | AZDS00000000 | NR_036789 |
| *Lactobacillus* sp. Sy-1 | JACCIU000000000 | MW715007 |

Table S7. Mean and standard deviation of ANI, AAI, and cAAI among the MAGs within each of the four observed groups and the other control groups (intra-clade ANI).

| Group | N. Genomes | ANI | AAI | cAAI |
|---|---|---|---|---|
| Group 1 | 3 | 99.72 ± 0.29 | 99.83 ± 0.15 | 99.89 ± 0.10 |
| Group 2 (-barcode20) | 6 | 99.71 ± 0.16 | 99.73 ± 0.14 | 99.93 ± 0.03 |
| Group 2 (+barcode20) | 7 | 96.08 ± 6.40 | 97.30 ± 4.27 | 98.68 ± 2.20 |
| Group 3 | 4 | 99.82 ± 0.10 | 99.82 ± 0.10 | 99.98 ±0.01 |
| Group 4 | 4 | 99.40 ± 0.42 | 99.55 ± 0.30 | 99.83 ± 0.01 |
| *Nicolliela* + L. Sy-1 | 3 | 87.11 ± 9.11 | 87.07 ± 9.22 | 93.05 ± 4.98 |
| *A. jinshanensis* | 3 | 91.62 ± 9.34 | 94.22 ± 6.42 | 95.99 ± 4.46 |

Table S8. Mean, standard deviation of ANI among the MAGs between each of the four observed groups with and without barcode20 (inter-clade ANI).

| Groups | ANI | AAI | cAAI |
|---|---|---|---|
| G1 vs G2(-barcode20) | 81.32 ± 2.52 | 70.87 ± 1.50 | 76.39 ± 0.01 |
| G1 vs G2(+barcode20) | 80.89 ± 2.56 | 70.65 ± 1.50 | 76.34 ± 0.14 |
| G1 vs G3 | NA | 64.35 ± 0.12 | 70.69 ±0.02 |
| G1 vs G4 | NA | 64.62 ± 0.11 | 71.07 ± 0.02 |
| G2(-barcode20) vs G3 | NA | 64.30 ±0.06 | 71.15 ± 0.01 |
| G2(+barcode20) vs G3 | NA | 64.32 ± 0.09 | 71.09 ±0.13 |
| G2(-barcode20) vs G4 | NA | 64.77 ± 0.05 | 71.68 ± 0.02 |
| G2(+barcode20) vs G4 | NA | 64.81 ± 0.10 | 71.63 ± 0.12 |
| G3 vs G4 | 80.68 ± 0.13 | 79.32 ± 0.10 | 85.10 ±0.01 |

*NA results correspond to values much less than 60, which are not calculated by the program

Table S9. Mean and standard deviation of ANI, AAI, and cAAI among the MAGs within each of the four observed groups vs the two *Nicoliellas* genomes.

| Group | N. Genomes | ANI vs closest relative | AAI vs *Nicoliella* | cAAI vs *Nicoliella* |
|---|---|---|---|---|
| Group 1 | 3 | 78.08 ± 0.09 *(Nicoliella)* | 68.83 ± 0.27 | 76.31 ± 0.15 |
| Group 2 (-barcode20) | 6 | 80.12 ± 0.41 *(Nicoliella)* | 79.35 ± 0.86 | 88.62 ± 0.65 |
| Group 2 (+barcode20) | 7 | 80.26 ± 0.52 (*Nicoliella*) | 79.42 ± 0.86 | 88.57 ± 0.66 |
| Group 3 | 4 | NA (*Acetilactobacillus*) | 64.35 ± 0.13 | 71.06 ±0.05 |
| Group 4 | 4 | NA (*Acetilactobacillus*) | 64.74 ± 0.26 | 71.53 ± 0.05 |

*NA results correspond to values much less than 60, which are not calculated by the program

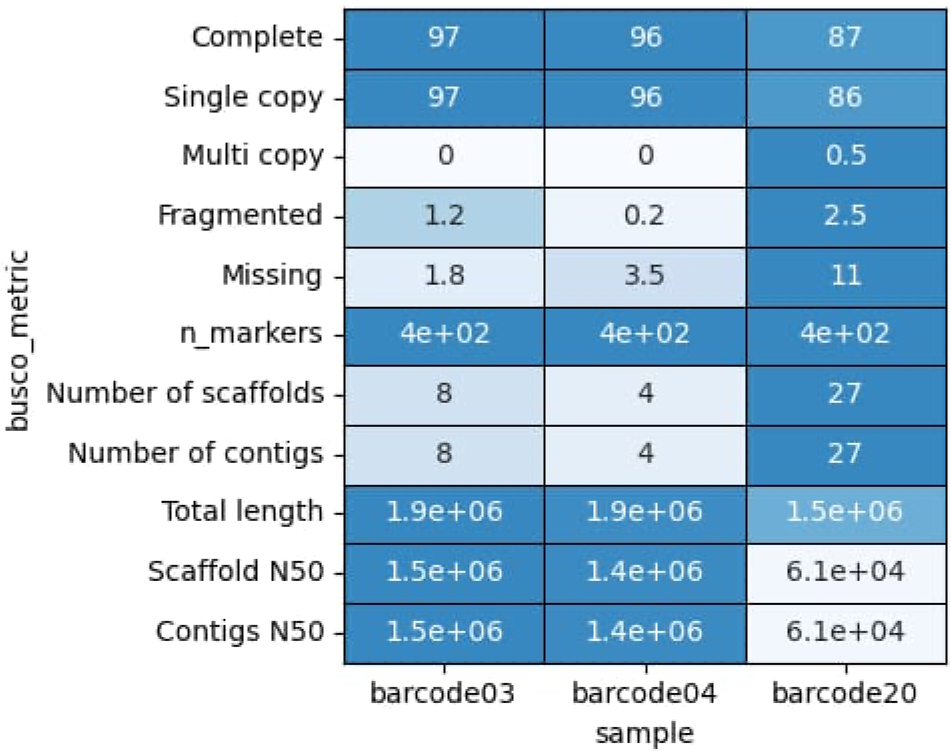
busco_metric
Complete
Single copy
Multi copy
Fragmented
Missing
n_markers
Number of scaffolds
Number of contigs
Total length
Scaffold N50
Contigs N50
97
96
87
97
96
86
0
0
0.5
1.2
0.2
2.5
1.8
3.5
11
4e+02
4e+02
4e+02
8
4
27
8
4
27
1.9e+06
1.9e+06
1.5e+06
1.5e+06
1.4e+06
6.1e+04
1.5e+06
1.4e+06
6.1e+04
barcode03
barcode04
barcode20
sample

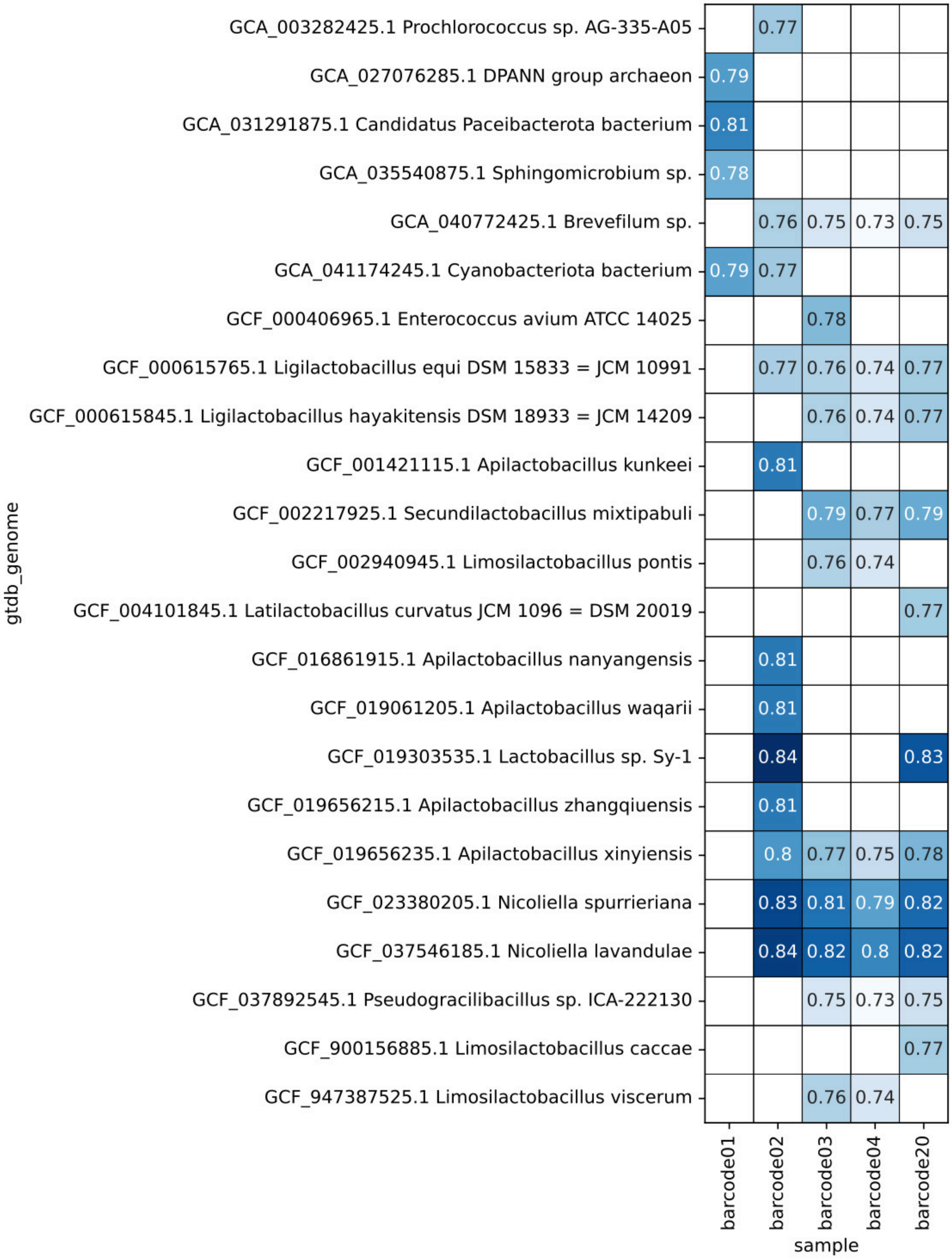
GCA_003282425.1 Prochlorococcus sp. AG-335-A05
GCA_027076285.1 DPANN group archaeon
GCA_031291875.1 Candidatus Paceibacterota bacterium
GCA_035540875.1 Sphingomicrobium sp.
GCA_040772425.1 Brevefilum sp.
GCA_041174245.1 Cyanobacteriota bacterium
GCF_000406965.1 Enterococcus avium ATCC 14025
GCF_000615765.1 Ligilactobacillus equi DSM 15833 = JCM 10991
GCF_000615845.1 Ligilactobacillus hayakitensis DSM 18933 = JCM 14209
GCF_001421115.1 Apilactobacillus kunkeei
GCF_002217925.1 Secundilactobacillus mixtipabuli
GCF_002940945.1 Limosilactobacillus pontis
GCF_004101845.1 Latilactobacillus curvatus JCM 1096 = DSM 20019
GCF_016861915.1 Apilactobacillus nanyangensis
GCF_019061205.1 Apilactobacillus waqarii
GCF_019303535.1 Lactobacillus sp. Sy-1
GCF_019656215.1 Apilactobacillus zhangqiuensis
GCF_019656235.1 Apilactobacillus xinyiensis
GCF_023380205.1 Nicoliella spurrieriana
GCF_037546185.1 Nicoliella lavandulae
GCF_037892545.1 Pseudogracilibacillus sp. ICA-222130
GCF_900156885.1 Limosilactobacillus caccae
GCF_947387525.1 Limosilactobacillus viscerum
gtdb_genome
0.77
0.79
0.81
0.78
0.76 0.75 0.73 0.75
0.79 0.77
0.78
0.77 0.76 0.74 0.77
0.76 0.74 0.77
0.81
0.79 0.77 0.79
0.76 0.74
0.77
0.81
0.81
0.84 0.83
0.81
0.8 0.77 0.75 0.78
0.83 0.81 0.79 0.82
0.84 0.82 0.8 0.82
0.75 0.73 0.75
0.77
0.76 0.74
barcode01
barcode02
barcode03
barcode04
barcode20
sample

Table S10. Reported *Acetilactobacillus* genus in the literature.

| **Taxon** | **Sample origin** | **Ref** | **Abundance** | **Collection date** | **Annotation date** | **Num samples** | **Technology** | **Size (Mb)** | **pH** |
|---|---|---|---|---|---|---|---|---|---|
| *A. jinshanensis* HSLZ-75 (CP034726) | Zhenjiang aromatic vinegar | (Yu *et al.*, 2020) | - | 2015 | 2018 | 1 | Illumina isolate strain | 1.6 | 4.0 |
| *Lactobacillus PRJNA379892* | Baijiu alcohol fermentation stage | (Liu P. & Miao, 2020) | *>95%* | 2020 | 2017 | 5 Baiyunbian baijiu fermentation batches | *16S rDNA amplicons* | - | - |
| *Lactobacillaceae* | Spoiled Sichuan bran vinegar | (Wang *et al.*, 2023) | >95% | 2020 | 2020 | 10 samples | *16S rDNA amplicons* | - | - |
| *A.jinshanensis subsp. aerogenes* Z-1 CP051512/CP051513 | Spoiled Sichuan bran vinegar | (Wang *et al.*, 2023) | - | 2020 | 2020 | 36 isolates | Illumina MiSeq and PacBio isolate strain | 1.6 +1 plasmid | 3.6 |
| *A. jinshanensis BJ01* PRJNA886850 | Baijiu fermented grains | (Liao *et al.*, 2023) | - | 2019 | 2022 | 9 lactic and/or acetic acid | RNA Illumina HiSeq 2500 | - | 3.5 |
| *A.jinshanensis* brasiliensis CP171343.1 | Supernatant from bacteria isolated from larval food of stingless bees | (Santos *et al.*, 2022) | | 2019 | 2024 | 1 MAG | BGISEQ | 1.1 | |
| Two clades of *Acetilactobacillus* PRJNA1111860 | *Mellipona* bees and honey samples. | (Cerqueira *et al.*, 2024) | more in bees than honey | 2017 | 2024 | 18 samples | 16S rRNA Illumina | - | - |
| *A. jinshanensis L214* PRJNA886850 CRA015377 CRA015377 and CRA013941 | Baijiu fermented grain | (Chen *et al.*, 2025) | 80% average on 28 day | - | - | 1 isolate | Transcriptomics, proteomics, and metabolomics | | |
| *A. jinshanensis* PRJCA026508 | Sichuan Baoning vinegar | (Wu J. *et al.*, 2025) | 23.02 % | 2022 | 2024 | 3 ponds * 11 time points=33 sequence | Illumina shotgun 150 bp not public | - | - |
| *A. jinshanensis* | Sichuan | | 50% | 2022 | 2024 | 3 ponds * | Illumina | - | - |

| | | | | | | | |
|---|---|---|---|---|---|---|---|
| PRJCA026508 | Baoning vinegar | (Liu A. *et al.*, 2025) | on 25 day | | | 11 time points=33 sequence | shotgun 150 bp not public |
